# Redefining De Novo Gammaherpesvirus Infection Through High-Dimensional, Single-Cell Analysis of Virus and Host

**DOI:** 10.1101/2020.08.11.203117

**Authors:** Jennifer N. Berger, Bridget Sanford, Abigail K. Kimball, Lauren M. Oko, Rachael E. Kaspar, Brian F. Niemeyer, Kenneth L. Jones, Eric T. Clambey, Linda F. van Dyk

**Affiliations:** Department of Immunology and Microbiology, University of Colorado Anschutz Medical Campus | Aurora, CO, 80045, USA; Department of Pediatric Oncology, University of Colorado Anschutz Medical Campus | Aurora, CO, 80045, USA; Department of Anesthesiology, University of Colorado Anschutz Medical Campus | Aurora, CO, 80045, USA; Department of Cell Biology, University of Oklahoma Health Sciences Center | Oklahoma City, OK, 73104, USA

## Abstract

Virus infection is frequently characterized using bulk cell populations. How these findings correspond to infection in individual cells remains unclear. Here, we integrate high-dimensional single-cell approaches to quantify viral and host RNA and protein expression signatures using de novo infection with a well-characterized model gammaherpesvirus. While infected cells demonstrated genome-wide transcription, individual cells revealed pronounced variation in gene expression, with only 9 of 80 annotated viral open reading frames uniformly expressed in all cells, and a 1000-fold variation in viral RNA expression between cells. Single-cell analysis further revealed positive and negative gene correlations, many uniquely present in a subset of cells. Beyond variation in viral gene expression, individual cells demonstrated a pronounced, dichotomous signature in host gene expression, revealed by measuring host RNA abundance and post-translational protein modifications. These studies provide a resource for the high-dimensional analysis of virus infection, and a conceptual framework to define virus infection as the sum of virus and host responses at the single-cell level.

**HIGHLIGHTS:**

- CyTOF and scRNA-seq identify wide variation in gene expression between infected cells.
- Host RNA expression and post-translational modifications stratify virus infection.
- Single cell RNA analysis reveals new relationships in viral gene expression.
- Simultaneous measurement of virus and host defines distinct infection states.

## INTRODUCTION

Gammaherpesviruses (γHVs) are a subfamily of the *Herpesviridae* characterized by their ability to establish latency in lymphoid cells and their association with numerous malignancies (Barton et al., 2011; Cesarman, 2014; Zamora, 2011). The γHVs include the human viruses Kaposi’s sarcoma-associated herpesvirus (KSHV, or human herpesvirus 8) and Epstein-Barr virus (EBV), and murine gammaherpesvirus 68 (MHV68 or γHV68; ICTV nomenclature, *murid herpesvirus 4*, MuHV-4), a model system for its human virus counterparts (Barton et al., 2011; Dong et al., 2017; Virgin et al., 1997). MHV68 provides a genetically tractable system, capable of inducing a range of outcomes *in vitro* and *in vivo* (Barton et al., 2011; Forrest and Speck, 2008; Suarez and van Dyk, 2008).

Like other herpesviruses, the MHV68 lifecycle is characterized by two major stages: latent and lytic infection (Pellett and Roizman, 2013). During latent infection, viral gene expression is limited, with no *de novo* viral replication (Pellett and Roizman, 2013). While latent γHV infection is associated with lifelong infection and malignancies, the γHVs critically rely on lytic infection for virus replication and transmission between hosts. Lytic infection is characterized by robust viral gene expression and the production of new infectious virions. The conventional view of lytic infection is that of a synchronous cascade of gene expression that results in genome-wide transcription. The paradigm for lytic gene expression was established in bulk cell populations, stratifying viral gene expression into three broad categories: immediate-early (IE), early (E), and late (L) gene expression (Johnson et al., 2010; Pellett and Roizman, 2013; Rochford et al., 2001). IE genes are expressed in the absence of new viral protein synthesis, E genes require new viral protein synthesis but do not require new viral DNA synthesis, and L genes require new viral DNA synthesis (Ahn et al., 2002; Ebrahimi et al., 2003; Johnson et al., 2010; Martinez-Guzman et al., 2003; Pellett and Roizman, 2013). Lytic gene expression can also be stratified by kinetic class, affording a revised perspective on viral gene expression (Cheng et al., 2012).

The advent of high-throughput, single-cell based methods has revealed a more complex picture of herpesvirus infection, from latent infection (Messinger et al., 2019; Oko et al., 2019; Shnayder et al., 2018), to lytic infection (Drayman et al., 2019; Oko et al., 2019; Wyler et al., 2019), reactivation from latency (Adang et al., 2006; Oko et al., 2019) and viral tropism (Sen et al., 2014). Recently, we identified unanticipated heterogeneity between infected cells during lytic infection, with wide variance in viral gene expression, from cells with robust viral gene expression to cells with little to no detectable expression of either viral non-coding RNAs or mRNAs (Oko et al., 2019). These studies suggest that lytic infection may be more heterogeneous in individual cells than the perspective revealed by analysis of bulk populations.

Though stages of infection are defined based on viral gene expression, lytic infection is also associated with profound alterations in host gene expression. First, many herpesviruses, including KSHV, EBV, MHV68, varicella zoster virus (VZV), and Herpes simplex virus (HSV), induce widespread degradation of host RNAs (i.e. host shutoff) during lytic infection, a phenomenon associated with decreased expression of cellular housekeeping genes in either bulk cell populations (Glaunsinger, 2015; Smiley, 2004; Waterboer et al., 2002) or individual cells (Oko et al., 2019). Viruses have also evolved diverse mechanisms to manipulate host processes (e.g. cell cycle, apoptosis, protein synthesis and signal transduction) (Chang et al., 2016). For example, the ORF36 gene of MHV68 and EBV encodes a viral kinase that phosphorylates H2AX, a proximal marker of the DNA damage response, to facilitate viral replication (Tarakanova et al., 2007). Recent studies have further demonstrated that the EBV and KSHV ORF36 homologs target additional host proteins including SAMHD1 and S6K to impact virus restriction and protein synthesis, respectively (Bhatt et al., 2016; Zhang et al., 2019). These studies emphasize that measuring host processes commonly targeted by multiple viruses [including host shutoff (Glaunsinger, 2015), DNA damage responses (Weitzman and Fradet-Turcotte, 2018) and innate signaling (Lee et al., 2015)], may provide an important, complementary strategy to define viral infection.

Here we provide a resource to redefine lytic infection from a single-cell perspective, using cytometry by time-of-flight (CyTOF, or mass cytometry) and single cell RNA-seq to achieve high-dimensional, single-cell analysis of viral and host gene expression. These studies reveal: i) the highly variable nature of viral gene expression between individual cells, with only a small subset of genes expressed across all cells, ii) the utility of measuring host parameters as a discriminator to identify infection state, and iii) new insights into infection and gene expression relationships, previously obscured by bulk cell analysis.

## RESULTS

### CyTOF analysis identifies a distinct protein expression profile for virally-infected cells

Recently, we discovered significant variability in viral gene expression between cells during MHV68 infection (Oko et al., 2019). These studies identified a prominent fraction of cells with limited viral gene expression, a phenomenon that could arise due to asynchronous infection, infection with restricted viral gene expression (e.g. latency), or cells resistant to infection. To define the complexity of virus infection in productively infected cells, we established a system to purify cells based on expression of the MHV68 latency-associated nuclear antigen (LANA) protein, a viral gene product expressed during lytic and latent infection (Rochford et al., 2001). NIH 3T12 fibroblasts, a highly permissive cell line for MHV68 lytic infection, were infected with a wild-type MHV68 recombinant expressing a LANA::β-lactamase gene fusion (WT MHV68.LANAβlac), followed by FACS purification (Diebel et al., 2015; Nealy et al., 2010) and high-dimensional single-cell analysis using CyTOF and single cell RNA-sequencing (scRNA-seq).

First, we analyzed virally-infected cells by CyTOF, to obtain a high-dimensional single-cell analysis of protein expression (Kimball et al., 2018). NIH 3T12 fibroblasts were harvested at 16 hours post-infection (hpi), a time at which no cytopathic effect or new progeny virus is released, and sort purified into LANAβlac+ (LANA+) and LANAβlac- (LANA-) fractions (Figure S1A). On average, >94% of cells were LANA+ at this time, indicating active virus infection in the vast majority of cells (Niemeyer et al., 2018). Mock-infected 3T12 cells were purified in parallel. To ensure robust cross-comparison between these populations, each sample was subjected to isotopic palladium-based barcoding, pooled together, stained with a panel of 23 isotopically-labelled antibodies, with samples collected on a Helios mass cytometer (Figure 1A). Mock-infected, LANA+ and LANA- cells had pronounced differences in clustering and protein expression profiles (Figure S1B-D). LANA+ cells robustly expressed the virally-encoded regulator of complement activation (vRCA) protein (Kapadia et al., 1999) and phosphorylated histone H2AX (pH2AX), a target of the ORF36 viral kinase (Tarakanova et al., 2007). LANA+ cells further showed increased expression of cell surface proteins (CD9, MHC I) and cell cycle proteins (Ki-67, phosphorylated Rb [Serine 807/S811]), with modest induction of signal transduction pathways (activated beta-catenin, pERK1/2, p-p38, pStat1, pStat3, pAKT and pS6) compared to both LANA- and mock-infected populations (Figure S1C-D). These data indicate that at the population level, LANA+ cells have a distinct protein expression signature relative to mock-infected and LANA- cells.

**Figure 1.**
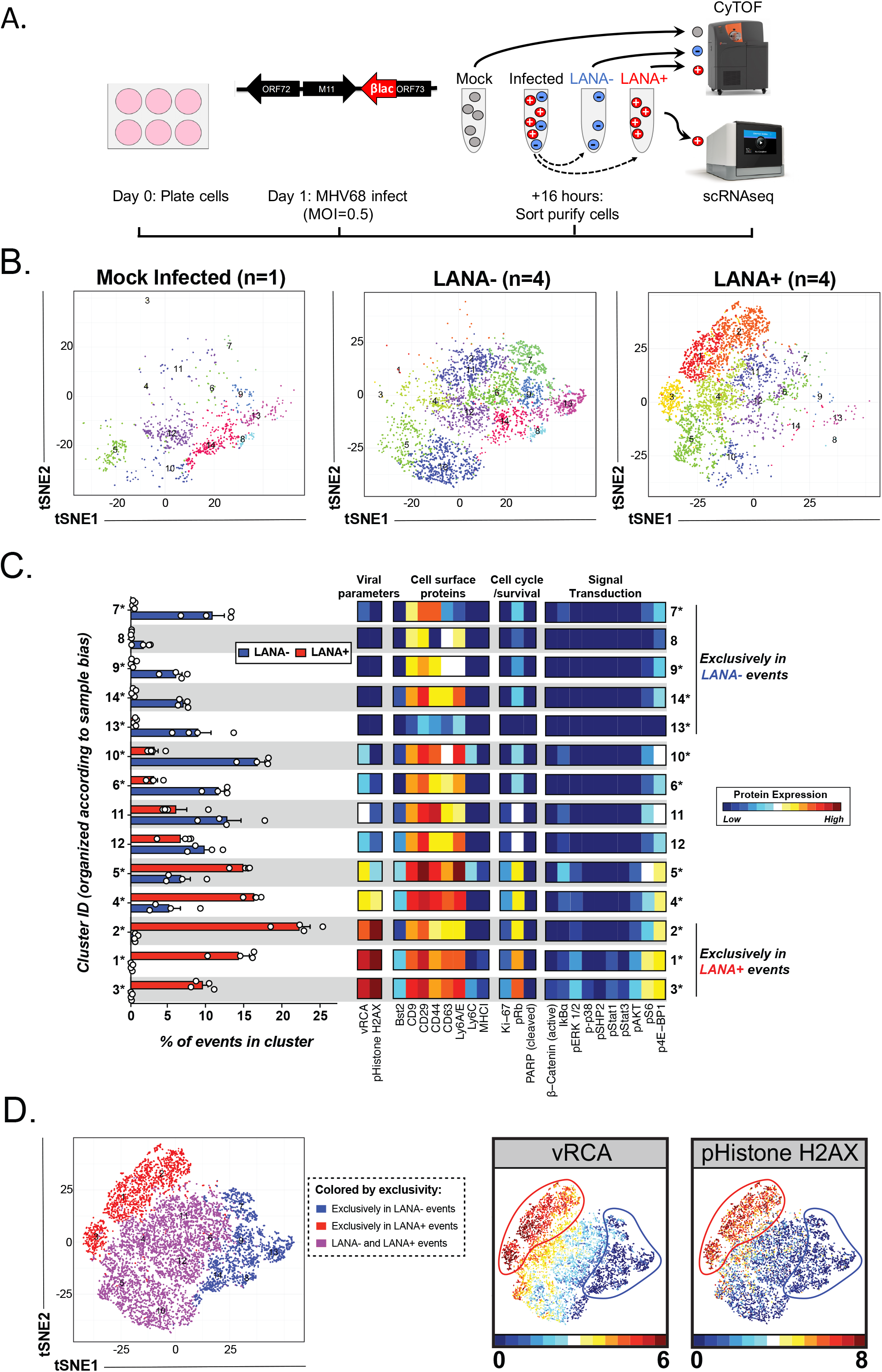
CyTOF analysis identifies a distinct protein expression profile for virally-infected cells. High-dimensional single-cell protein analysis of MHV68 infection by CyTOF. (A) 3T12 fibroblasts were either mock or MHV68-infected with WT MHV68.LANAβlac (MOI=0.5), with indicated populations FACS purified according to LANA::β-lactamase expression at 16 hours pi. (B) PhenoGraph analysis of mock, LANA- and LANA+ cells subjected to CyTOF analysis and visualized by tSNE-based dimensionality reduction identified 14 cell clusters (colored by cluster ID). (C) Comparison of the frequency (left) and protein expression profile (right) of PhenoGraph-defined clusters in LANA- and LANA+ samples. Clusters (in rows), ranked from those exclusively in LANA- samples (top) to those exclusively in LANA+ samples (bottom); asterisks indicate clusters statistically significantly different between conditions, defined by unpaired t-test corrected for multiple comparisons using the Holm-Sidak method, p<0.05. Data show mean ± SEM with individual symbols indicating individual sample values (n=4 per group). Relative protein expression for each cluster (right panel) is indicated by heatmap intensity, with protein markers divded by functional categories. (D) tSNE plot of LANA- and LANA+ cells colored according to LANA- and LANA+ exclusivity (left), vRCA expression (middle), and pH2AX expression (right panel). Red and blue lines identify events exclusive to LANA+ or LANA- clusters (left panel). Ruler defines range of expression with values calculated using the equation arcsinh (x/5) where x is raw expression value. Live, DNA+ cells (^191^Ir+ ^193^Ir+ ^195^Pt-) were imported into PhenoGraph with 9,162 events total analyzed (1,018 events from each sample; n=1, mock, n=4 each for LANA+ and LANA- samples), clustered on 9 cell surface proteins (vRCA, BST2, CD9, CD29, CD44, CD63, Ly6A/E, Ly6C, and MHCI). See also Figure S1.

Next we quantified the diversity of cellular phenotypes between LANA+ and LANA- cells. When we applied PhenoGraph, an unsupervised clustering algorithm (Levine et al., 2015), we identified 14 phenotypes across LANA+, LANA-, and mock-infected populations (Figure 1B). The frequency of phenotypic clusters varied widely between LANA+ and LANA- populations (Figure 1C), with 3 clusters exclusively present among LANA+ cells, 5 clusters uniquely present among LANA- cells, and 6 clusters containing both LANA+ and LANA- cells (Figure 1C). Clusters exclusively present among LANA+ cells (clusters 1-3, at the bottom of Figure 1C) were characterized by maximal expression of vRCA and pH2AX, and increased expression of pRB, pERK1/2, pAKT, and pS6 relative to other cell clusters (Figure 1C). Expression of other parameters, including CD9, CD29, CD44, CD63, Ki-67 and IkBα did not strictly correlate with either LANA+ or LANA- cells, indicating that these proteins were not solely regulated as a function of virus infection (Figure 1C). While high expression of vRCA and pH2AX was a strong discriminator between clusters exclusively in LANA+ versus LANA- events, approximately half of LANA+ events expressed low levels of either vRCA or pH2AX, overlapping with LANA- cells (Figure 1C-D). These data indicate that while many cells expressing the LANAβlac fusion protein had increased virus and host protein expression, there was a substantial fraction of LANA+ cells that could not be readily distinguished from LANA- cells (clusters colored purple, Figure 1D). These data further identify phosphorylation of H2AX (i.e. pH2AX) as a robust marker of virus-induced phenotypic changes at the single-cell level.

### LANA+ cells show heterogeneous protein and post-translational modifications during lytic infection

To better understand phenotypic heterogeneity among cells with active viral gene expression (i.e. LANA+ cells), we quantified phenotypic diversity among a large number of LANA+ cells, clustering cells based on expression of nine cell surface-expressed proteins (BST2, CD9, CD29, CD44, CD63, Ly6A/E, Ly6C, MHCI, and vRCA). This focused clustering analysis identified 15 phenotypic clusters within LANA+ cells, which were then visually inspected based on vRCA, pH2AX, Ly6C, and BST2 expression (Figure 2A-C). LANA+ cells could be subdivided into cells with high and low viral gene expression, with vRCA and pH2AX expression tightly overlapping. In contrast, Ly6C and BST2 protein expression was variable across vRCA^high^ pH2AX^high^ and vRCA^low^ pH2AX^low^ cell fractions, indicating that these proteins did not simply correlate with virus infection (Figure 2B-C). Stratification of LANA+ cells into pH2AX^high^ and pH2AX^low^ subsets (Figure S2A) demonstrated that pH2AX^high^ cells had increased expression of vRCA, MHC class I, and multiple phosphorylation-dependent modifications including pRb, active β-catenin, pERK1/2, pStat1 and pAKT compared to pH2AX^low^ cells (Figure 2D). pH2AX^high^ cells also had modestly decreased expression of CD9, CD29, Ly6A/E, Ly6C and IkBα relative to pH2AX^low^ events (Figure 2D). Consistent with the tight correlation between vRCA and pH2AX expression, LANA+ vRCA^high^ cells had increased expression of pH2AX, MHC class I, pRb, active β-catenin, pERK1/2, pStat1 and pAKT compared to vRCA^low^ cells and modestly reduced IkBa expression (Figure S2B-E). These studies indicate that vRCA and pH2AX are strong markers of progressive infection among LANA+ events, with phosphorylated Rb a robust secondary marker associated with a pH2AX^high^ vRCA^high^ phenotype. These results further indicate that heterogeneous progression through infection can be revealed by monitoring host protein expression at the single-cell level.

**Figure 2.**
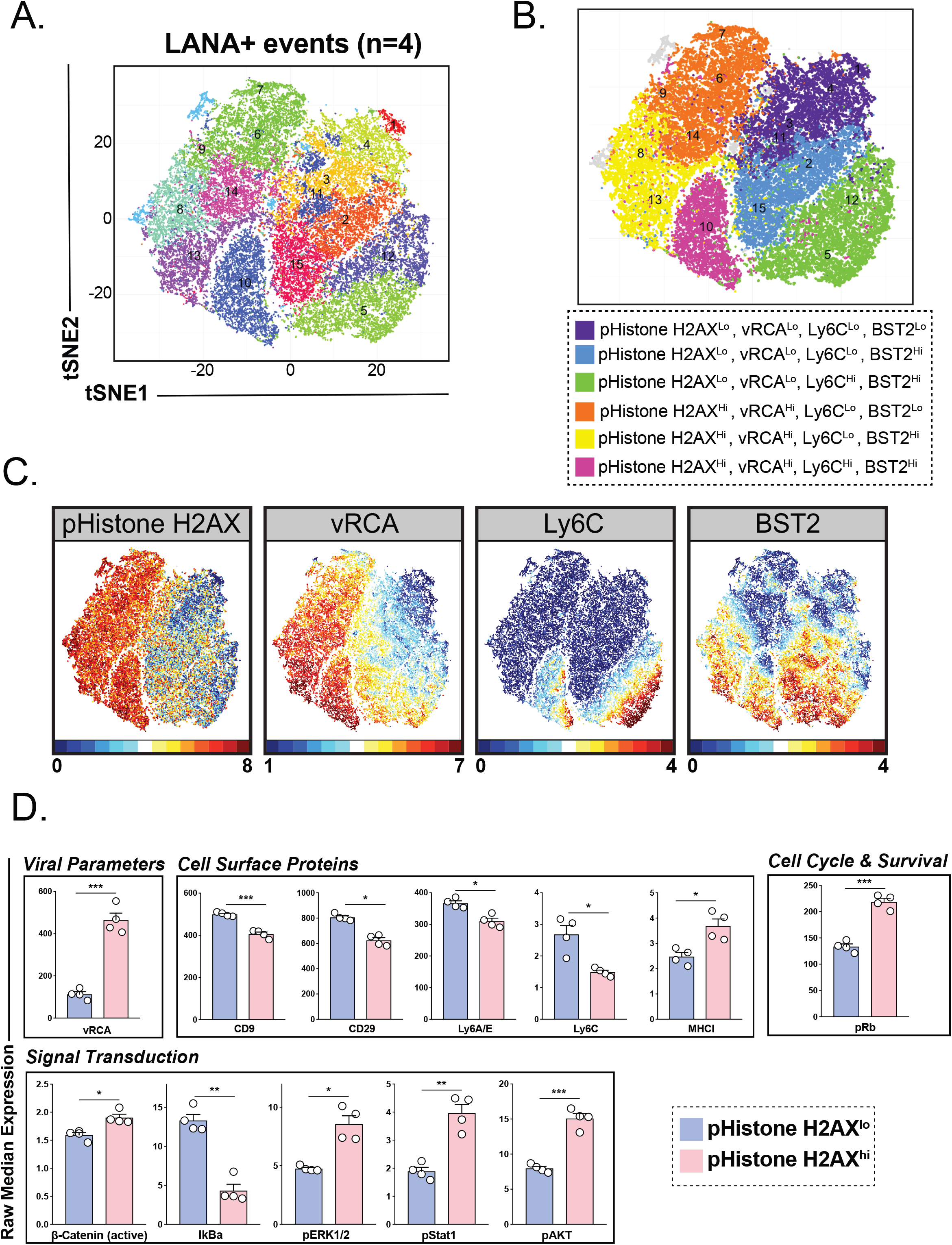
LANA+ cells show heterogeneous protein and post-translational modifications during lytic infection. CyTOF analysis of LANA+ MHV68-infected cells as in Figure 1. (A) PhenoGraph analysis of 40,000 LANA+ cells (10,000 cells/sample, 4 samples) clustered by 9 cell surface proteins (vRCA, BST2, CD9, CD29, CD44, CD63, Ly6A/E, Ly6C, and MHCI) identified 15 unique cellular clusters colored by cluster ID and displayed on a tSNE plot. (B) LANA+ cells can be stratified into six categories based on expression of pH2AX, vRCA, Ly6C, and BST2, with (C) LANA+ cells colored by expression levels for each protein marker. Ruler defines range of expression with values calculated using the equation arcsinh (x/5) where x is raw expression value. (D) LANA+ cells were divided into pH2AX^Lo^ and pH2AX^Hi^ events, and compared for raw median expression. Data depict proteins whose expression was statistically significantly different between pH2AX^Lo^ and pH2AX^Hi^ events. Data depict mean ± SEM with individual symbols indicating values from independent samples (n=4/group). All samples were analyzed for statistical significance using unpaired t tests, corrected for multiple comparisons using the Holm-Sidak method, with statistical significance denoted as *p<0.05, ** p<0.01, ***p<0.001. See also Figure S2.

### MHV68-infected cells with a pH2AX^high^ vRCA^high^ cell phenotype express high viral mRNA and low host actin RNA

We next investigated how pH2AX and vRCA expression corresponded with viral and host RNA expression at the single-cell level, using the PrimeFlow methodology to simultaneously measure protein and RNAs (Oko et al., 2019). This analysis focused on two RNAs which we previously reported showed heterogeneous expression during MHV68 infection (Oko et al., 2019): i) ORF18, a viral RNA transcribed with late kinetics and ii) Actb, a host RNA encoding beta-actin that is known to be targeted for degradation by the MHV68 SOX endonuclease (Covarrubias et al., 2009).

As expected, mock-infected cells had no vRCA expression and a low frequency of pH2AX+ events (Figure 3A). In contrast, MHV68-infected 3T12 cultures exhibited multiple populations, including: 1) pH2AX- vRCA- cells (population 1, gray box, Figure 3A bottom panel), 2) pH2AX+ vRCA- cells (population 2, light blue box), 3) pH2AX- vRCA+ cells (population 3, orange box), and 4) pH2AX+ vRCA+ cells (population 4, red box). As these studies did not use purified LANA+ cells, in contrast to the CyTOF studies, the presence of these four populations suggests that these cultures contained a mixture of infected and uninfected cells.

**Figure 3.**
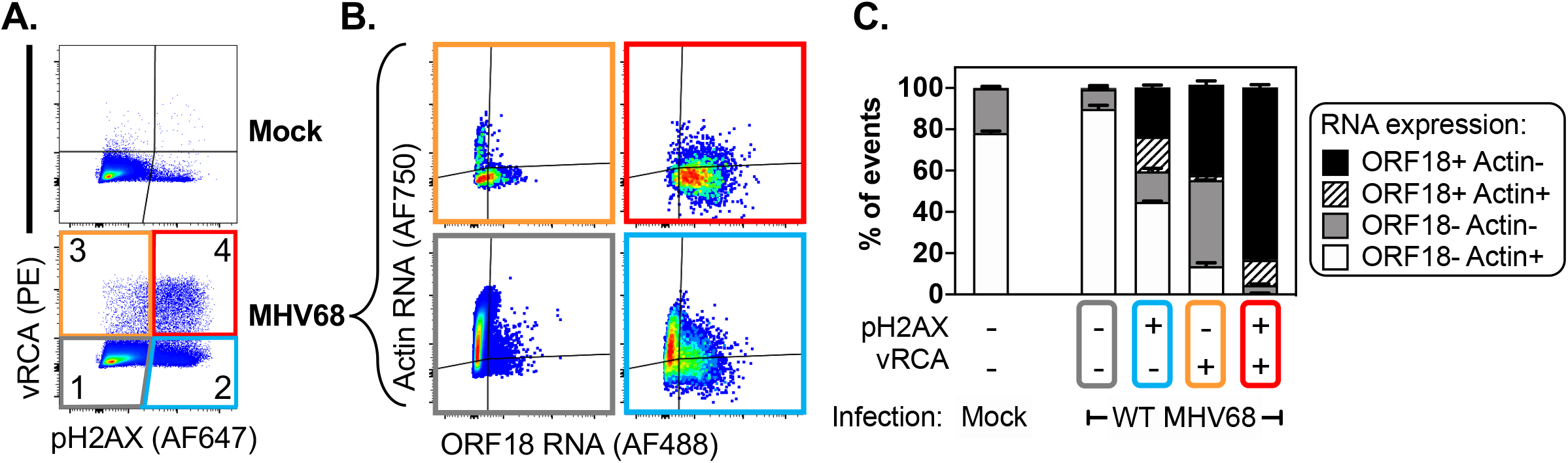
MHV68-infected cells with a pH2AX^high^ vRCA^high^ cell phenotype express high viral mRNA and low host actin RNA. Flow cytometric analysis of protein and RNA expression in 3T12 fibroblasts infected with MHV68 (MOI=0.5, harvested 16 hpi). (A) Analysis of vRCA and pH2AX protein expression in mock and MHV68-infected cultures identified four populations denoted by numbers and color (1=Gray, 2=Blue, 3=Orange, 4=Red). (B) Analysis of viral ORF18 and host Actin (Actb) mRNA expression across four populations of MHV68-infected cultures, stratified by vRCA and pH2AX expression. Border color of biaxial plots corresponds to the parent population stratified by protein expression in panel A. (C) Quantitaiton of the frequency of cells based on RNA expression profile as in panel B, depicting mean ± SEM from 3 independent experimental samples. Flow cytometric analysis was done on singlets, following gating to remove doublets.

We next queried RNA expression within cells subdivided by pH2AX and vRCA expression, using ORF18 expression as a direct readout of virus infection and Actb degradation as a secondary readout of virus infection. Cells with a pH2AX- vRCA- phenotype (gray box, Figure 3A-B) were present in mock- and MHV68-infected cultures, with the vast majority of cells characterized by an ORF18- Actb+ phenotype and no evidence of infection (Figure 3B-C). In contrast, cells with a pH2AX+ vRCA- phenotype (light blue box, Figure 3A-B) contained ~1/3 of cells that were ORF18+ with variable expression of Actb (Figure 3B-C), identifying infection based on viral RNA expression in the absence of a viral protein. Cells with a pH2AX- vRCA+ phenotype (orange box, Figure 3A-B) were dominated by ORF18+ Actb- and ORF18- Actb- cells, demonstrating direct measures of infection by RNA and protein. Lastly, pH2AX+ vRCA+ cells (red box, Figure 3A-B) were predominantly ORF18+ Actb-, identifying viral RNA and viral protein expression in the context of host RNA degradation. These data demonstrate that pH2AX+ vRCA+ cells show progressive infection at the RNA and protein level, whereas cells expressing only vRCA or pH2AX represent intermediate infection phenotypes as measured by viral RNA expression and host RNA degradation.

### scRNA-seq reveals wide variation in the frequency and magnitude of viral RNA expression

We next sought to obtain a global perspective on how MHV68 infection affects gene expression using scRNA-seq. 3T12 fibroblasts were infected with an MOI of 0.5 PFU/cell using two viruses with indistinguishable lytic replication [WT MHV68.LANAβlac and a viral cyclin-deficient (cycKO) MHV68.LANAβlac (van Dyk et al., 2000)], with LANA+ cells sort purified at 16 hpi (Figure 1A). LANA+ cells were submitted for bulk RNA-seq and scRNA-seq analysis, with a mean of 94,810 reads per cell for scRNA-seq (Figure S3A-C). While there was some concordance between the most abundant host RNAs observed in bulk RNA-seq and scRNA-seq, scRNA-seq analysis revealed wide variability among cells, a level of complexity obscured in bulk RNA-seq analysis (Figure S3D). We next focused on scRNA-seq, given its capacity to define intercellular variation and to avoid averaging of signal intensity across a population of cells. Comparison of scRNA-seq data between WT and CycKO infected cells demonstrated a similar distribution of cells and gene expression, with only five genes with average gene expression differences greater than 2-fold (Table S1 and S2), consistent with their comparable lytic replication in vitro (van Dyk et al., 2000). Therefore, subsequent analysis integrated scRNA-seq data from LANA+ cells purified from WT and CycKO infection.

To understand MHV68 transcription at the single-cell level, we first quantified the diversity and magnitude of viral transcripts. When viral RNAs (enumerated as unique molecular identifiers, UMIs) were mapped to the MHV68 genome, 77 of the 80 annotated MHV68 ORFs were detected. M10b, M10c and M12, three viral RNAs derived from MHV68 repeat structures were the only viral RNAs not detected (Virgin et al., 1997). LANA+ cells expressed a wide range of viral genes (12-66 viral genes) per cell, with a median of 52 viral genes per cell, based on detection of ≥ 1 UMI per viral gene per cell (Figure 4A). Mean UMI per viral gene and percentage of cells expressing viral genes were positively correlated (Spearman r coefficient = 0.9775), with 86.2% of LANA+ cells expressing the ORF73 RNA (Figure 4B). Despite cell purification based on uniform LANA expression, total viral UMIs per cell spanned a >1000-fold range (19-24,718 viral UMIs per cell) (Figure 4C). While the number of viral genes detected per cell was positively correlated with total viral UMIs per cell, LANA+ cells exhibited a bimodal distribution of virus RNA^high^ (Virus^high^) cells (>500 viral UMIs/cell) and virus RNA^low^ (Virus^low^) cells (<500 viral UMIs/cell) (Figure 4C-D). The distinction between Virus^high^ and Virus^low^ cells was not simply explained by how many viral genes were detected per cell, as some cells expressed the same number of viral genes yet showed a >10-fold variance in total viral UMIs (e.g. the range of viral UMIs among cells that express 40 viral genes per cell, Figure 4D).

**Figure 4.**
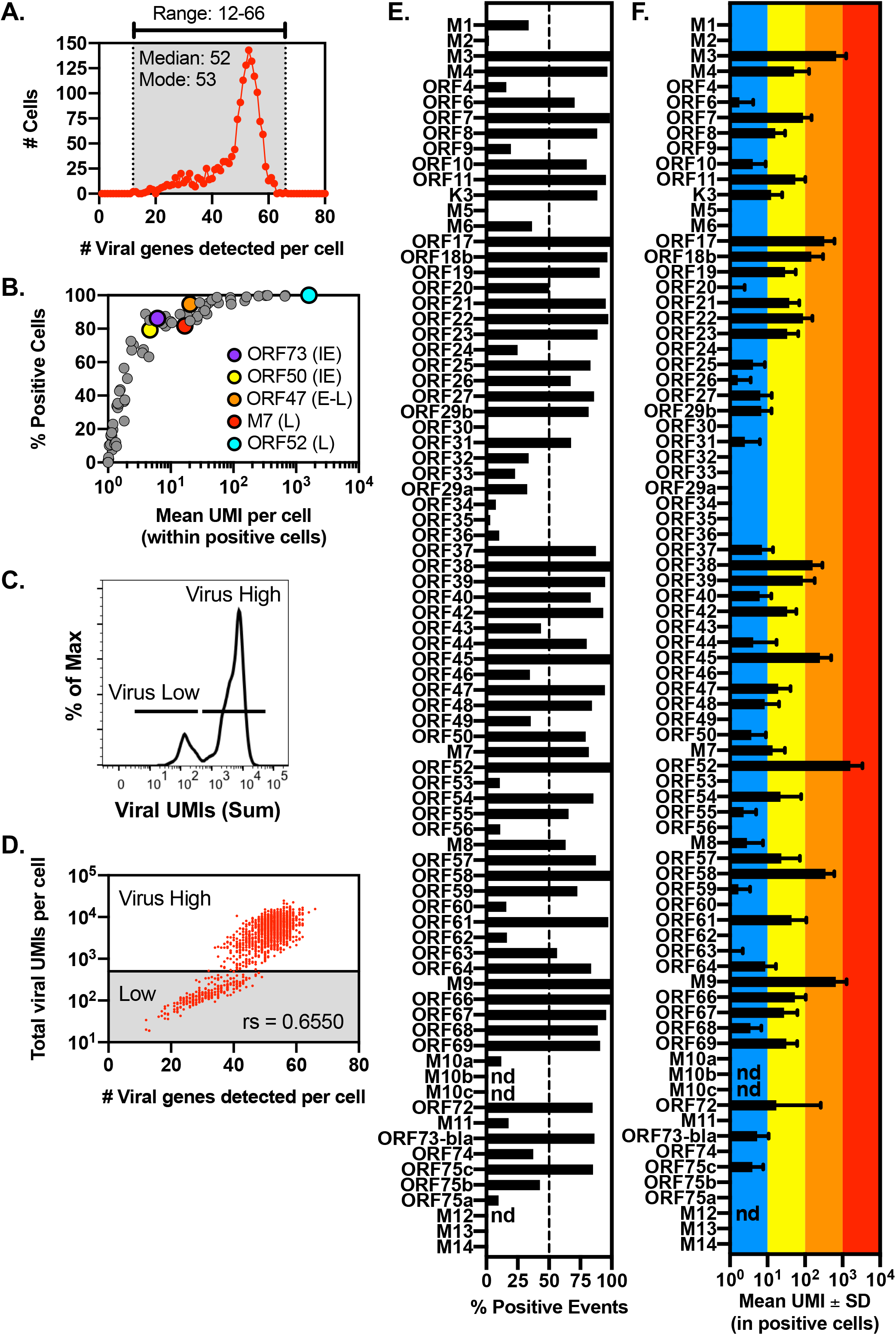
scRNA-seq reveals wide variation in the frequency and magnitude of viral RNA expression. scRNA-seq analysis of MHV68 viral gene expression in LANA+ 3T12 fibroblasts as in Figure 1A. (A) LANA+ cells show a wide range in how many viral genes are expressed on a per-cell basis. Expression of a viral gene was defined conservatively as any viral gene with ≥1 UMI detected per cell, with data showing the distribution of cells based on number of viral genes expressed per cell. (B) Viral genes vary widely in mean UMI per cell and the frequency of cells expressing an individual viral gene. Mean viral UMI per cell was calculated from positive cells that expressed ≥1 UMI per gene, with five representative genes identified by unique shading. Each dot represents expression for a single viral gene product for all 80 annotated MHV68 open reading frames. (C) The distribution of cells based on total viral UMIs reveals a bimodal distribution of virus low and virus high cells. (D) Total viral UMIs per cell as a function of the number of viral genes detected per cell (as in panel A). Cells demonstrate a bimodal distribution of total viral UMIs with virus high and low cells, with a positive Spearman correlation coefficient, rs, as indicated. (E) Frequency of cells expressing each viral gene, with viral gene expression defined as ≥1 UMI for each viral gene. Dashed line indicates 50% of events. (F) Mean UMI count per viral gene, defined by calculating mean value only from positive cells (≥1 UMI for each viral gene), depicting mean ± SD. Data represent scRNA-seq data from all LANA+ cells (n=1605 cells). nd, genes that were not detected. See also Figure S3 and Table S1.

To better understand viral transcription at the single-cell level, we quantified the percentage of cells expressing each viral gene (Figure 4E) and mean UMI count within positive cells (Figure 4F). The percentage of cells which expressed viral genes varied widely, with 9 viral genes (M3, ORF7, ORF17, ORF38, ORF45, ORF52, ORF58, M9, and ORF66) expressed in the vast majority (98.63-100%) of cells (Figure 4E). These ubiquitous viral RNAs spanned transcriptional categories [IE (ORF38), Early-Late (M3, ORF7), and L genes (ORF17, ORF45, ORF52, ORF58, M9, ORF66)] and kinetic classes [class I (ORF38), class II (M3, ORF45, ORF52, ORF58), class III (ORF17, M9, ORF66) and class IV (ORF7)] (Cheng et al., 2012). In contrast, many viral genes were only detected in a subset of cells, including ORF50, an IE gene that encodes Rta, a major transcriptional regulator of lytic replication (Figure 4E). The range of expression for individual genes varied widely (Figure 4F). While many viral genes had a mean UMI per viral gene between 1-10 UMIs per cell, the most highly expressed viral gene ORF52 had a mean of >1000 UMIs per cell.

### Visualization of genome-wide viral gene expression at the single-cell level

To investigate the complexity of gene expression across MHV68-infected cells, we next visualized the absolute UMI count for each viral gene across all 1,605 LANA+ cells. Cells were rank-ordered based on total viral UMIs detected per cell, from Virus^low^ to Virus^high^ cells across a 1000-fold range of viral UMIs per cell, with each row depicting expression for each MHV68 viral gene and each column depicting data from a single cell (Figure 5). This continuum of gene expression revealed multiple patterns of gene expression. First, there were some viral genes (e.g. ORF59 and ORF73βlac, which encodes LANA) which were detected in almost all cells, with only modest differences in expression between cells with the fewest viral UMIs and the greatest viral UMIs (Figure 5). Second, viral genes such as M3 and ORF52 progressively increased in expression, proportional to the increase in viral UMIs (Figure 5). Third, viral genes, including ORF48, ORF54, and ORF57, showed maximal expression in cells with intermediate viral counts (Figure 5). By stratifying cells into either Virus^high^ or Virus^low^ categories (a bifurcation identified in Figure 4C-D), we identified a subset of genes including K3, ORF42 and ORF44 whose expression was sporadic in Virus^low^ cells and uniform in Virus^high^ cells (Figure 5 and Figure S4). ORF68 was one of the only genes whose expression was slightly higher among Virus^low^ cells compared to Virus^high^ cells (Figure 5 and Figure S4). Gene expression signatures for Virus^low^ and Virus^high^ cells did not clearly track with any transcriptional or kinetic class, and individual genes within these distinctions demonstrated significant gene to gene variation (Figure S5A). These data identify the highly variable nature of MHV68 transcription between individual cells during de novo infection.

**Figure 5.**
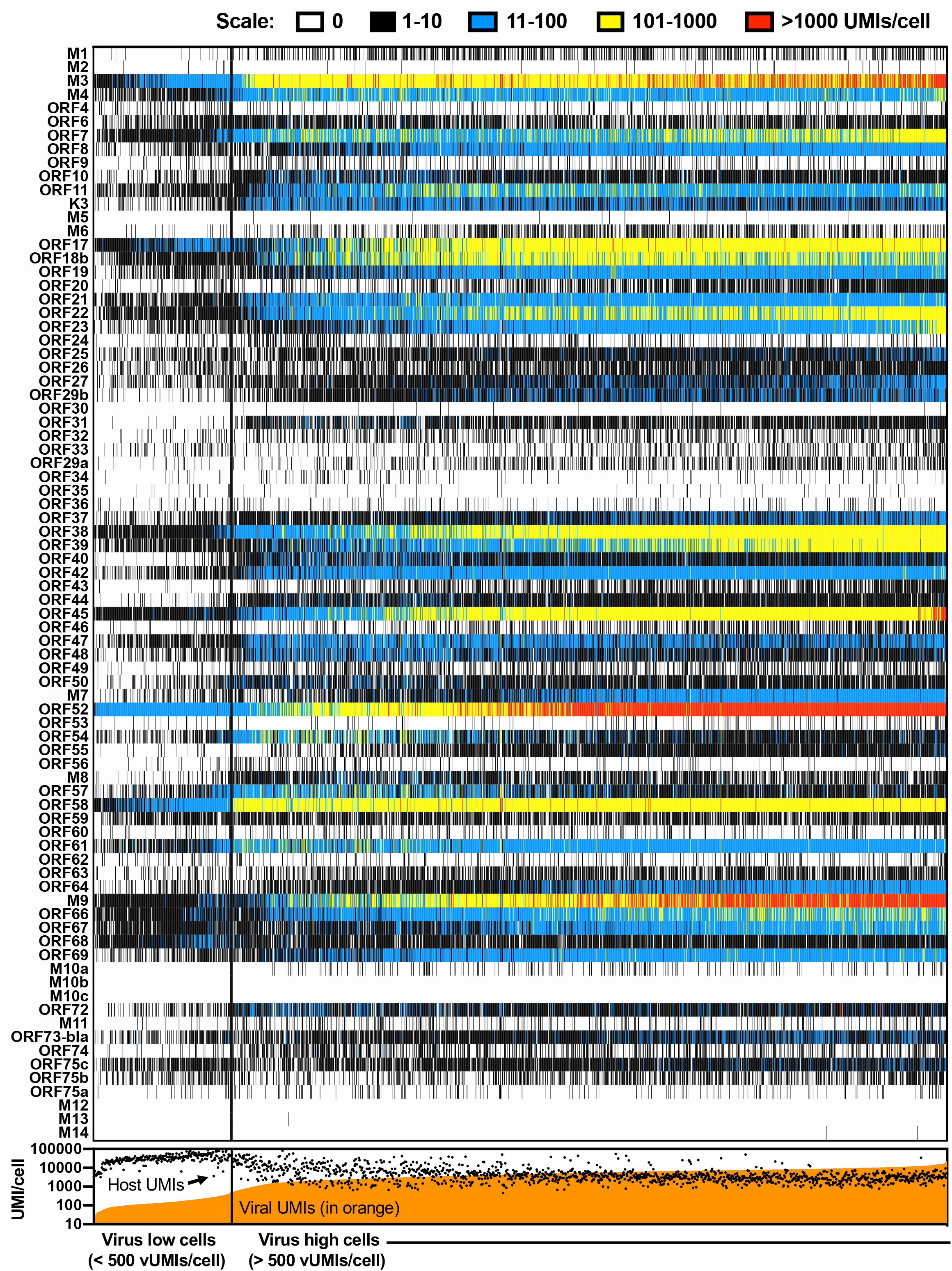
Visualization of genome-wide viral gene expression at the single-cell level. scRNA-seq analysis of MHV68 gene expression in LANA+ 3T12 fibroblasts as depicted in Figure 1A. Comprehensive visualization of viral gene expression on a single cell basis. Each row depicts expression pattern for a different viral gene. Each column depicts expression within an individual cell. Cells are rank ordered from the cell with the least (left) to the greatest total viral UMI count (right), with corresponding total viral (filled orange line) and host (black dots) UMIs per cell depicted in the bottom panel. Cells are further bisected into Virus low cells (< 500 viral UMIs per cell) and Virus high cells (> 500 viral UMIs per cell) by a thick vertical black line, from a total of 1605 cells. UMI counts are stratified based on the indicated heatmap scale. See also Figure S4.

### MHV68-infected cells can be stratified by differential abundance of virus and host RNAs

To clarify the patterns of gene expression that we observed, we used unsupervised K-means clustering to identify related cells. By clustering on viral and cellular RNA expression, we identified 5 clusters among LANA+ cells (Figure 6A). 3-dimensional tSNE-based data visualization further identified that an IE gene (ORF50) had maximal expression in clusters A and B, an early-late (E-L) gene (ORF47) had maximal expression in clusters B and C, and a L gene (M7) had maximal expression in clusters D and E (Figure 6B and Figure S5B). In contrast, host Actb RNA, a target of virus-induced host shutoff mediated by the ORF37 gene, showed maximal expression in clusters A and B, with low expression across clusters C through E (Figure 6B-C). These data suggested that at this timepoint, LANA+ cells had heterogeneous virus-induced host shutoff. Cells with a high percentage of viral UMIs were predominant in clusters C through E, in contrast to cells with a high frequency of host UMIs prevalent in clusters A and B (Figure 6D). Based on these distinctions, we divided cells into virus-based and host-biased groups (Figure 6E-F).

**Figure 6:**
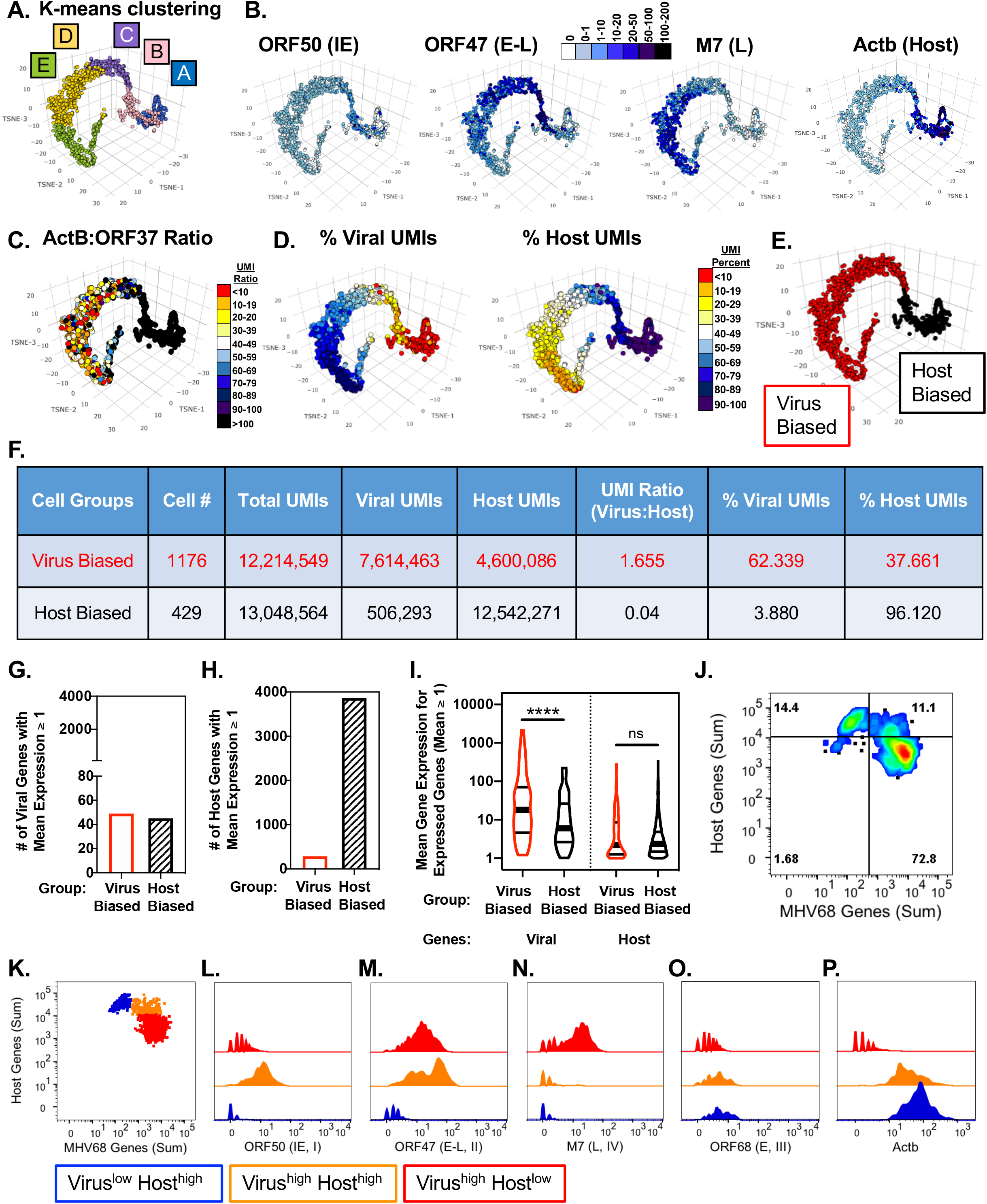
MHV68-infected cells can be stratified by differential abundance of virus and host RNAs. scRNA-seq analysis of MHV68-infected LANA+ cells for viral and host RNAs. (A-E) 3D tSNE-based dimensionality reduction was used to visualize gene expression within individual cells. (A) K-means clustering identified 5 unique cell clusters, as labelled. (B) Viral and host RNAs show different patterns of expression, comparing viral immediate early (IE), early-late (E-L) and late (L) genes with host beta actin (Actb). UMI counts visualized in individual cells (symbols) as defined by key. (C) The ratio of host Actb to ORF37 UMIs, calculated for each cell and overlaid onto tSNE-based visualization, varies widely across cells. (D) Percent of viral (left) and host UMIs (right) per cell was calculated for each cell, portrayed by shade of color overlaid onto tSNE visualization. (E) Cells were separated into two groups: virus biased (red) and host biased (black) based on percent viral and host UMIs, and K-means clustering. Host biased cells included clusters A and B with <20% virus UMIs on a per cell basis. (F) Characteristics of virus and host biased cells based on scRNA-seq data. (G-H) Number of (G) viral or (H) host genes with mean UMI expression ≥1, among either virus (open red symbol) or host biased cells (dashed black line). (I) Violin plots depict the distribution of mean gene expression among expressed genes (mean UMI expression ≥1), comparing viral genes (left) or host genes (right) in either virus biased (open red symbol) or host biased cells (open black line). Quartiles and median are depicted in each violin plot. Statistical comparisons compared viral or host genes and their difference between virus and host biased cells. \*\*\*\**p*<0.0001, one way ANOVA with Tukey’s multiple comparisons test. ns, not significant. (J-K) Biaxial analysis of LANA+ cells (n=1605 cells) comparing total host UMI count (y axis) versus total viral UMI count (x axis) identified three major cell populations, stratified by differential expression of viral or host genes. (K) Identification of cells that are Virus^low^ Host^high^ (blue), Virus^high^ Host^high^ (orange), and Virus^high^ Host^low^ (red). (L-P) Histogram overlays comparing relative expression of (L-O) viral and (P) host genes for the three identified cell populations. Each gene is denoted by its IE, E, or L class and by its kinetic class defined by (Cheng et al., 2012). See also Figure S5-6, Table S2.

The different proportions of viral and host UMIs between virus- and host-biased cells could result from changes in either how many viral or host genes were expressed, or mean expression per gene. Notably, the number of virus genes expressed (defined by mean UMI per cell ≥ 1) was comparable between virus-biased and host-biased cells (Figure 6G). In contrast, virus-biased cells were characterized by a pronounced decrease in the number of host RNAs expressed, compared to host-biased cells (Figure 6H). When we analyzed gene expression levels, virus-biased cells had increased viral gene expression per cell compared to host-biased cells, with no significant difference in host gene expression among expressed genes (Figure 6I). These data indicate that virus-biased cells have increased viral gene expression, coupled with a pronounced reduction in the number of detectable host RNAs. While cell cycle-associated gene signatures were relatively comparable between virus-biased and host-biased cells, host-biased cells expressed a number of interferon-response genes (Figure S6 and Table S3), suggesting a potential role in resistance to infection.

Although stages of virus infection are typically defined by viral parameters only, the observations of: i) Virus^high^ and Virus^low^ cells (Figure 4D), and ii) Virus-biased and host-biased cells (Figure 6G), suggested that these two processes may be connected. Indeed, when we analyzed the distribution of cells based on the sum of all host UMIs versus the sum of all viral UMIs, we found three prominent populations (Figure 6J-K): 1) 72.8% of cells had high viral RNA expression with reduced host RNA expression (i.e. Virus^high^ Host^low^, in red, Figure 6K), 2) 11.1% of cells had a Virus^high^ Host^high^ phenotype (in orange, Figure 6K), and 3) 14.4% of cells had a Virus^low^ Host^high^ phenotype (in blue, Figure 6K). The number of Virus^low^ Host^high^ cells almost completely corresponded with the number of Virus^low^ events which we previously identified (Figure 4D), with less than two percent of cells expressing a Virus^low^ Host^low^ phenotype (Figure 6J). By comparing viral gene expression across these 3 populations, we found that an IE gene (ORF50) and E-L gene (ORF47) had peak expression in Virus^high^ Host^high^ cells, intermediate expression in Virus^high^ Host^low^ cells, with limited expression in Virus^low^ Host^high^ cells (Figure 6L-M). In contrast, a L gene (M7) had high expression exclusively in Virus^high^ Host^low^ cells (Figure 6N). Virus^low^ Host^high^ cells had lower expression across many virus genes, with the exception of ORF68 which was modestly higher in Virus^low^ Host^high^ cells compared to other cells (Figure 6O). Actb, a known target of virus-induced shutoff, retained high expression in both Virus^low^ Host^high^ and Virus^high^ Host^high^ cells with a pronounced decrease in Virus^high^ Host^low^ cells (Figure 6P). These data demonstrate that while many LANA+ cells show signs of robust viral RNA expression coupled with reduced host RNAs, subsets of LANA+ cells express high levels of host RNAs with differential viral RNA abundance.

### Correlation analysis reveals positive and negative correlations in MHV68 transcription at the single-cell level

Finally, we sought to investigate the inter-relationships between viral genes, leveraging the unique insights obtained from single-cell data. Initially, we generated a correlation matrix of all viral gene-gene expression relationships querying all LANA+ cells. This correlation matrix identified that the majority of gene-gene pairs had a positive correlation (83.6%, depicted in blue), with fewer negative correlations (16.1%, depicted in red) (Figure S7). We next examined gene-gene relationships in Virus^high^ or Virus^low^ cells. While the majority of gene-gene relationships in Virus^high^ and Virus^low^ cells were positive correlations, Virus^high^ cells revealed an increased frequency of negative correlations (29.1%) among a subset of viral genes (Figure 7A and S7).

**Figure 7.**
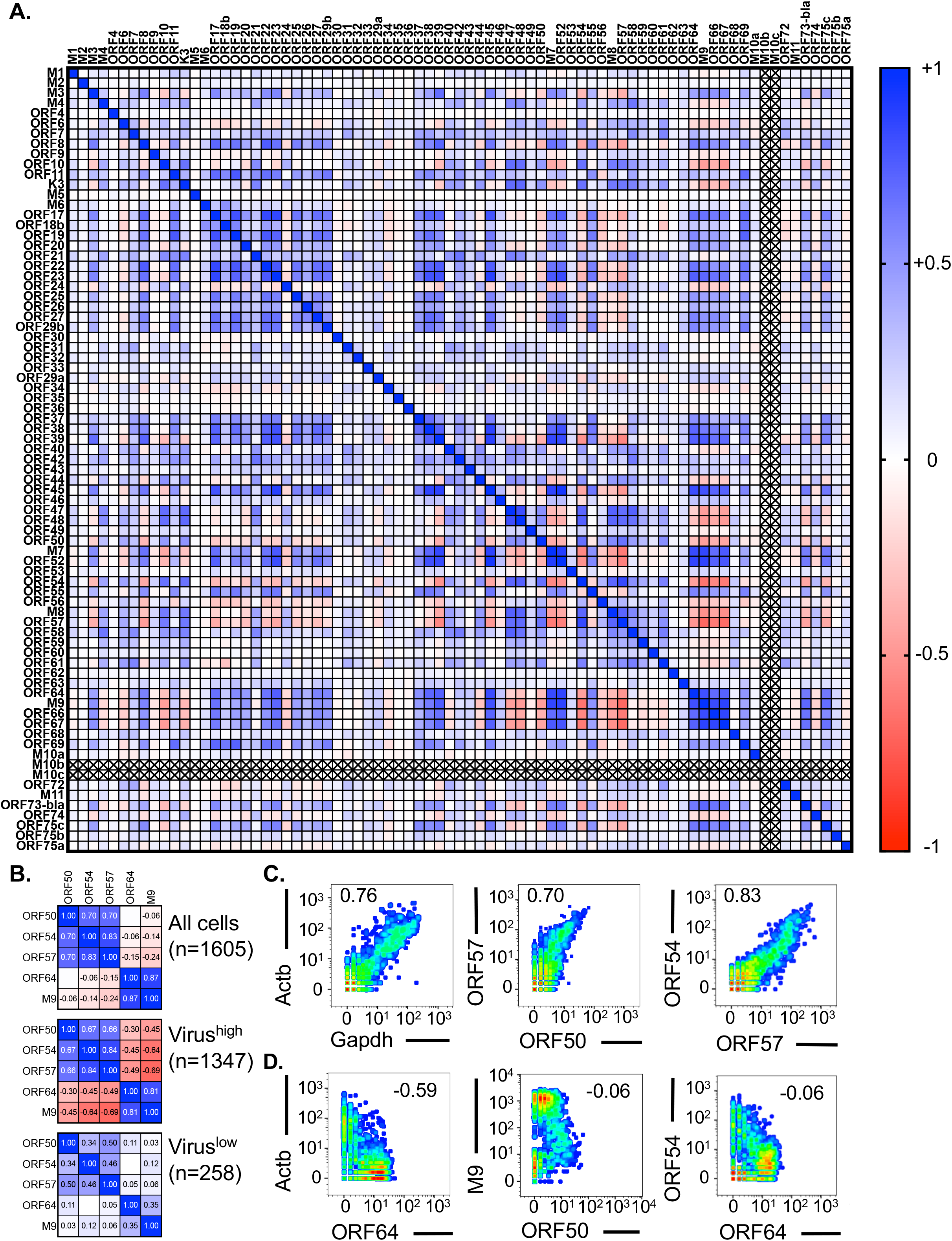
Correlation analysis reveals positive and negative correlations in MHV68 transcription at the single-cell level. The inter-relationship of viral gene expression was interrogated using scRNA-seq data from MHV68-infected LANA+ cells. (A) Correlation matrix of viral genes among Virus^high^ cells (>500 viral UMIs/cell), depicting Spearman correlation coefficient, rs, for each gene pair according to the indicated heatmap. Positive correlation is indicated by blue, negative correlation indicated by red. Correlation matrix depicts all viral genes except for M10b and M10c (X indicates gene for which there were no values) and M12, M13 and M14 (excluded due to negligible values). (B) A comparison of correlation coefficients between viral gene pairs when analyzing all cells (top), Virus^high^ (middle) or Virus^low^ (bottom) cells. Data identify discrepant correlations observed in different cell subsets. (C) Examples of host and virus gene pairs with positive correlations, demonstrated by plotting scRNA-seq gene expression values on biaxial plots across all LANA+ cells. (D) Examples of host and virus gene pairs with a lack of correlation or negative correlation. Panels in C-D depict scRNA-seq values for all LANA+ cells (n=1,605 cells), including both Virus^high^ and Virus^low^ events. Each plot includes the Spearman correlation coefficient for the indicated gene pair defined across all cells. See also Figure S7.

We next sought to investigate gene-gene relationships as of function of Virus^high^ or Virus^low^ cells. On the one hand, there were some gene pairs whose relationship was preserved between Virus^high^ and Virus^low^ cells, exemplified by ORF50 and ORF57, and ORF54 and ORF57, both showing positive correlations across Virus^high^ and Virus^low^ cells (Figure 7B-C). On the other hand, there were other gene pairs whose relationship was distinct between Virus^high^ and Virus^low^ cells. These included ORF50 and M9, and ORF54 and ORF64, with minimal correlation across total cells, a negative correlation in Virus^high^ cells, and minimal correlation among Virus^low^ cells (Figure 7B). The complexity of gene-gene expression relationships was clearly demonstrated by plotting gene-by-gene expression levels from total cells (Figure 7C-D). Predicted correlations were exemplified by a positive correlation between host RNAs Actb and Gapdh and a negative correlation between Actb and ORF64 (Figure 7C-D). While positive correlations between ORF50 and ORF57, and ORF57 and ORF54 were consistent across all events, the relationships between ORF50 and M9, ORF64 and ORF54 were more complex (Figure 7D). These data emphasize that gene-gene coexpression relationships can be obscured by combining cell subsets, and emphasize the value of single-cell approaches to define gene expression inter-relationships within homogeneous cell populations.

## DISCUSSION

In this manuscript, we present a strategy to investigate viral infection, simultaneously quantifying viral and host RNA and protein expression in individual cells. This approach provides a conceptual framework to understand how individual cells respond to virus infection at the protein and RNA level, presents new insights into the complexity of virus infection from virus and host perspective, and establishes high-dimensional datasets as a resource for further analysis of virus-host interactions at the single-cell level. By focusing on a well-characterized virus, tightly controlled infection conditions, and cells with uniform viral gene expression, these studies emphasize the knowable complexity of infection in individual cells and the limitations of studying infection in bulk cell populations.

### Implications

Our studies illustrate the power of defining virus infection by quantifying viral and host gene expression across individual cells and have significant implications for understanding virus-host interactions. First, our data highlight that gene expression analysis in a population of cells conceals wide variance in virus infection, specifically failing to identify cells with low gene expression and obscuring negative feedback mechanisms. Second, our data emphasize that defining virus infection strictly by viral gene expression ignores the essential contribution of the host cell to this process, and that virus infection stage may be more appropriately considered as the integrated impact of virus on the host. Quantitation of host RNAs and virus-dependent post-translational host modifications were particularly informative in discriminating between infection subtypes, while measures of protein abundance were less impactful. Third, our data suggest that phenotypic variation between cells, or input virions, results in cell subsets with restricted viral gene expression, a potentially novel strategy to identify negative regulators of infection. While the relative contribution of host and virus variation to these diverse outcomes remains to be addressed, there is an increasing breadth of literature in support of phenotypic variation from both sides (e.g. (Avraham et al., 2015; Chaturvedi et al., 2020; Hansen et al., 2018; Shaffer et al., 2017)). We propose that integrated single-cell analysis of virus and host will be particularly informative to understand how viral and host genes regulate infection, whether influencing viral or host gene expression, and/or altering the relative proportion of permissive or resistant cells. Fourth, these studies provide an opportunity to reassess fundamental concepts in virology, including how variation in virus (e.g. differing ratios in particle:PFU ratio (Van Skike et al., 2018)) or the host cell integrate to impact infection outcome. We anticipate that further studies, combined with further mining of these datasets by the community, will identify new rules governing virus infection that may be pharmacologically targeted.

### Insights into herpesvirus infection

Herpesvirus infection is classically divided into lytic infection, that generates new infectious virions, and latent infection, that facilitates long-term persistence in host cells while avoiding immune detection. Lytic infection is frequently understood as a coordinated cascade of gene expression, characterized by genome-wide transcription (Pellett and Roizman, 2013). High resolution mapping of coding and non-coding viral RNAs (Cheng et al., 2012; O’Grady et al., 2019) and new molecular insights into the regulation of viral RNA transcription and abundance (Hartenian et al., 2020) have further refined models of lytic gene expression. Despite these advances, most studies have analyzed bulk cell populations, obscuring intercellular heterogeneity.

Here, we provide three new datasets, with two complementary high-dimensional single-cell approaches, CyTOF and scRNAseq, to investigate *de novo γ*HV infection of cells that are highly permissive for lytic infection. These studies build on our recent analysis of MHV68 gene expression during *de novo* infection, studies that revealed asynchronous or inefficient infection among MHV68-exposed cells (Oko et al., 2019). Here, we extended these studies, purifying actively-infected cells (defined by expression of the viral protein LANA, which is expressed during lytic and latent infection) and deeply characterizing RNA and protein expression profiles during *de novo* infection. By monitoring viral and host RNAs and proteins, these studies defined robust markers of productive lytic infection and revealed heterogeneity in progression following *de novo* infection.

These studies initially focused on how MHV68 infection influences cellular responses at the protein level using CyTOF. By comparing cells with or without signs of active infection, we found multiple infection phenotypes. First, LANA+ cells contained a prominent cell fraction that co-expressed high levels of a viral protein, vRCA, and phosphorylation of histone H2AX, a virus-dependent event. vRCA^high^ pH2AX^high^ cells were only found in LANA+ cells, and were further characterized by increased phosphorylation of host proteins (pRb, active β-catenin, pERK1/2, pStat1 and pAKT) consistent with reports of γHV-induced modulation of the host proteome (Chang et al., 2016). vRCA^high^ pH2AX^high^ cells also had a prominent RNA phenotype, ORF18^high^ (late viral RNA) and Actb^low^ (host RNA indicator of virus-induced host shutoff). While vRCA^high^ pH2AX^high^ ORF18 RNA^high^ Actb RNA^low^ cells showed the most progressive signs of infection, we readily identified additional cell subsets expressing viral RNA in the absence of viral protein or host-shutoff, potentially reflecting delayed infection or a disconnect between transcription and translation. These studies identify vRCA and pH2AX as strong indicators of robust and progressive virus infection, and indicate that including additional viral and host indicators can distinguish among diverse infection stages.

We next turned to scRNA-seq to gain a broader understanding of virus and host RNA expression during *de novo* infection. Though we anticipated that many LANA+ cells (i.e. vRCA^high^ pH2AX^high^ cells) would express high viral RNAs and low host RNAs (exemplified by ORF18 and Actb, respectively), the relative homogeneity of LANA+ cells at the RNA level was unknown. One of the most striking outcomes of these studies was the wide range of viral and host gene expression observed at the single-cell level, derived from infection with a low MOI, and harvested at a single timepoint. LANA+ cells had a 1000-fold variance in total viral reads per cell, with cells expressing between 12-66 viral genes per cell and widespread variance in both the frequency and expression level of viral RNAs. While this analysis mapped RNAs to 77 of the 80 annotated MHV68 ORFs (Virgin et al., 1997) using a 3’ based sequencing platform, it is notable that: i) only one viral gene (ORF52) was detected in 100% of LANA+ cells, with ii) only 9 viral genes (M3, ORF7, ORF17, ORF38, ORF45, ORF52, ORF58, M9, and ORF66) expressed in a high percentage (>98.5%) of LANA+ cells. Though many of these genes were highly expressed on a per cell basis, a high frequency of expression does not solely track with high expression. For example, ORF73 RNA, which encodes for LANA, was detected in 86.2% of cells with an average of 5 UMIs per cell. Lack of detection of ORF73 in LANA+ sort purified cells may seem to be a paradox, as cells were purified based on LANA+ cells. This apparent discordance may either be due to RNA dropout or to cells expressing LANA protein in the absence of ORF73 RNA. For other genes, limited detection likely reflects extremely low expression during lytic infection [e.g. M2, a latency-associated gene that has not historically been detected during lytic infection (Rochford et al., 2001; Virgin et al., 1999)]. These results suggest that lack of detection of some MHV68 genes is likely due to little to no expression during lytic infection and not due to technical issues with scRNA-seq. More broadly, these studies indicate that stratifying gene expression by the IE→E→L paradigm, or by kinetic class, is insufficient to describe the complexity of viral transcription in individual cells, even in a highly permissive cell type.

An important complementary perspective on virus infection was our capacity to assess how MHV68 infection remodels host RNA expression. Virus-biased cells expressed 1.5 times more viral UMIs than host UMIs, a striking result when considering that there are only 80 annotated viral ORFs (Virgin et al., 1997) compared to ~25,000 host genes (Guenet, 2005). Virus:host RNA skewing was further driven by a dramatic reduction in host gene expression, consistent with virus-induced host shutoff (Covarrubias et al., 2011). Host shutoff was readily visualized by pronounced reduction in ActB RNA expression in vRCA^high^ pH2AX^high^ cells, illustrating the value of monitoring host RNAs at the single-cell level as an indicator of virus infection. When we assessed the global relationship between virus and host RNAs, we found additional complexity. While most LANA+ cells (73%) showed a reciprocal relationship expression of high viral RNAs and low host RNAs (i.e. Virus^high^ Host^low^), there were substantial cell fractions that were either Host^high^Virus^high^ or Virus^low^ Host^high^, suggesting delayed or impaired virus-induced host shutoff (a process that requires transcription, translation and nuclease-dependent degradation of RNAs (Covarrubias et al., 2011)).

### Potential relationships of virus infection phenotypes

The identification of distinct infection phenotypes raises the question of how these subsets are inter-related. Though definitive lineage relationships require sequential tracking of individual infected cells, we postulate that the Virus^high^ Host^high^ phenotype identifies cells with slower progression through lytic infection, preceding Virus^high^ Host^low^ cells. In contrast, we speculate that Virus^low^ Host^high^ cells may represent a distinct biological outcome. Although Virus^low^ cells expressed fewer viral genes, frequently at a lower magnitude, these cells contained viral RNAs from all phases of infection, with 90% of Virus^low^ cells expressing IE (ORF38), E or E-L (M3, ORF7, ORF68) or L (ORF17, ORF45, ORF52, ORF58, M9, ORF66) genes; these genes span kinetic classes, indicating that there is not a specific block in infection (Cheng et al., 2012). Virus^low^ cells did contain a few viral genes (e.g. ORF68) that are modestly increased relative to Virus^high^ cells; nonetheless, the Virus^low^ gene signature would be obscured in bulk cell analysis. The molecular basis of the Virus^low^ Host^high^ phenotype remains to be determined but could arise from host or virus variation. Curiously, Virus^low^ cells expressed multiple interferon-stimulated genes, but not type I interferons, suggesting a potential cell-intrinsic block to lytic infection. The Virus^low^ Host^high^ phenotype may also arise from virus-intrinsic differences (e.g. variation in particle:PFU differences (Klasse, 2015) or virus-like vesicles (Gong et al., 2017)). Variable outcomes of γHV infection have been reported in multiple contexts. MHV68 infection of endothelial cells results in a significant cell fraction of cells undergoing persistent infection (Suarez and van Dyk, 2008). KSHV infection of fibroblasts results in variable lytic or latent infection (Jang et al., 2016; Krishnan et al., 2004; Matthews et al., 2011), and EBV infection in gastric epithelial cells includes both lytic and latent outcomes (Hill et al., 2013; Ma et al., 2011; Nawandar et al., 2015). Despite the restricted gene expression of Virus^low^ cells, expression of multiple viral genes strongly suggests that these cells have the potential to impact biological outcomes and immune regulation.

### Virus gene-gene relationships revealed by scRNA-seq

Virus infection involves the coordination of highly complex gene expression patterns, a poorly understand process at the single-cell level. Given this knowledge gap, we leveraged the unique insights of scRNA-seq to quantify genome-wide correlation matrices. Unexpectedly, we found gene pairs which were differentially regulated between Virus^high^ or Virus^low^ subsets, including pronounced negative correlations among Virus^high^ cells. The complexity of these relationships was illustrated by examining correlations between ORF50 and other viral genes. ORF50 encodes the Rta viral transactivator and is known to both induce and repress transcriptional targets (Coleman et al., 2005; Damania et al., 2004; Hair et al., 2007; Pavlova et al., 2005; Wu et al., 2000). Correlation analysis among Virus^high^ cells corroborated established ORF50 targets (e.g. positive correlation with ORF57, negative correlation with ORF73) and identified new potential relationships (e.g. positive correlation with ORF54 and M8 and negative correlation with ORF64-67). These findings emphasize the importance of defining gene expression relationships in biologically homogeneous populations, something uniquely afforded by single-cell analysis.

Our scRNA-seq analysis quantified viral RNA abundance in LANA+ cells, mapping RNAs to the annotated MHV68 genome using polyA-primed sequencing. This resource illustrates the complexity of viral RNA expression following *de novo γ*HV infection and sets the foundation for new insights in gene expression in individual cells, including how virion-packaged RNAs shape expression, the relative contribution of transcription and RNA decay to the transcriptome, and RNA:protein relationships in individual cells (Hartenian and Glaunsinger, 2019; Purushothaman et al., 2015; Ruiz et al., 2019). This approach further emphasizes that not all abundant viral RNAs are equally expressed at the single-cell level, and argue for the importance of single-cell analysis to identify candidate immunogens based on uniform, high expression.

In total, these studies provide a resource to understand the complexity of virus-host interactions within individual cells. This work provides three databases from matched *de novo* gammaherpesvirus infection of a single permissive cell type, from CyTOF to bulk RNA-seq to scRNA-seq, and exemplify the power of these approaches to reveal new insights into virus-host interaction during γHV infection. We anticipate that these data will serve as an important benchmark for future studies of virus infection, providing a wealth of observations to further revise our understanding of virus-host interactions.

## ACKNOWLEDGEMENTS

The authors would like to acknowledge insightful comments made by the members of the Clambey and van Dyk laboratories, and the laboratories of Dr. Kelly Doran and Dr. Laurel Lenz for equipment and reagents. We acknowledge flow cytometry support through ClinImmune and the Department of Immunology & Microbiology, CyTOF services through the Flow Cytometry Shared Resource (FCSR) of the University of Colorado Cancer Center (UCCC), expert technical assistance regarding CyTOF from Christine Childs and Karen Helm, receipt of CyTOF antibodies through the coordinated efforts of the FCSR and the Human Immune Monitoring Shared Resource, and support of sequencing through the Genomics Shared Resource of the UCCC.

## AUTHOR CONTRIBUTIONS

<u>Conceptualization:</u> Jennifer N. Berger, Brian F. Niemeyer, Eric T. Clambey, Linda F. van Dyk

<u>Data curation:</u> Bridget Sanford, Abigail K. Kimball, Kenneth L. Jones, Eric T. Clambey

<u>Formal analysis:</u> Jennifer N. Berger, Bridget Sanford, Abigail K. Kimball, Kenneth L. Jones, Eric T. Clambey

<u>Funding acquisition:</u> Eric T. Clambey, Linda F. van Dyk

<u>Investigation:</u> Jennifer N. Berger, Lauren M. Oko, Rachael E. Kaspar, Brian F. Niemeyer

<u>Methodology:</u> Jennifer N. Berger, Bridget Sanford, Lauren M. Oko, Abigail K. Kimball, Rachael E. Kaspar, Brian F. Niemeyer, Kenneth L. Jones

<u>Project administration:</u> Eric T. Clambey, Linda F. van Dyk

<u>Resources:</u> Jennifer N. Berger, Bridget Sanford, Abigail K. Kimball, Kenneth L. Jones, Eric T. Clambey, Linda F. van Dyk

<u>Software:</u> Abigail K. Kimball, Eric T. Clambey, Bridget Sanford, Kenneth L. Jones

<u>Supervision:</u> Eric T. Clambey, Linda F. van Dyk

<u>Validation:</u> Jennifer N. Berger, Bridget Sanford, Abigail K. Kimball, Eric T. Clambey, Linda F. van Dyk

<u>Visualization:</u> Jennifer N. Berger, Bridget Sanford, Abigail K. Kimball, Kenneth L. Jones, Eric T. Clambey, Linda F. van Dyk

<u>Writing – original draft:</u> Jennifer N. Berger, Eric T. Clambey, Linda F. van Dyk

<u>Writing – review & editing:</u>

## STAR ⋆ METHODS

### Contact for Reagent and Resource Sharing

Further information and requests for resources and reagents should be directed to, and will be fulfilled by, the Lead Contact, Linda van Dyk.

### Experimental Model and Subject Details

#### Viruses

All experiments used WT MHV68 (strain WUMS; ATCC VR-1465) (Virgin et al., 1997), using either bacterial artificial chromosome–derived WT MHV68 (Adler et al., 2000) or WT MHV68.LANAβlac, which encodes a fusion between ORF73 and the beta-lactamase gene (Nealy et al., 2010). Certain studies were done using a variant engineered to contain a translational stop in the ORF72 (v-cyclin) gene, on the WT MHV68.LANAβlac background (Niemeyer et al., 2018). MHV68 was grown and titrated by plaque assay on 3T12 fibroblasts, as previously published (Diebel et al., 2015).

#### Cell Culture

NIH 3T12 cells (ATCC # CCL-164) were grown in Dulbecco’s modified Eagle medium (DMEM, Gibco, Life Technologies) supplemented with 5% fetal bovine serum (Atlanta Biologicals), 1x Penicillin-Streptomycin-L-Glutamine (Gibco, Life Technologies). Cells were passaged using 0.05% trypsin (Gibco, Life Technologies) after washing with 1x PBS (Gibco, Life Technologies). Cells were maintained at 37°C in a 5% CO_2_ humidified incubator (Forma Scientific).

### Method Details

#### RNA sequencing and Analysis

3T12 fibroblasts were infected at an MOI=0.5 with a low volume inoculum for one hour, after which inoculum was removed and cells were rinsed. Cells were harvested at 16 hours post-infection (hpi) and stained with β-lactamase substrate as previously reported (Diebel et al., 2015) using the CCF2-AM substrate (Invitrogen, Life Technologies). Cells were subjected to fluorescence activated cell sorting (FACS), with cells gated on singlet cells that were positive for the cleaved CCF2-AM substrate. Cleavage of CCF2-AM was defined as positive fluorescence in the violet laser-excited 450nm emission channel obtained in WT MHV68.LANAβlac or cycKO MHV68.LANAβlac-infected cultures, compared to parental WT MHV68 infected cultures with no beta-lactamase fusion gene, used to define background fluorescence. β-lactamase positive cells were sorted using a FACSAria Fusion (BD Biosciences). Cells were spun down at 1,200 RPM (Sorvall RT60003), and resuspended in 1x PBS (Gibco, Life Technologies). Cells were either prepared for CyTOF or RNA-sequencing by the Genomics and Microarray Shared Resource Core at University of Colorado. Bulk RNA-seq libraries were sequenced using Illumina HiSeq4000 platform with single-end reads (1 × 151), with forty million reads per sample collected and resulting sequences filtered and trimmed to remove low-quality bases (Phred score <15), and analyzed using a custom computational pipeline consisting of open-source gSNAP, Cufflinks. Single cell RNA Sequencing (scRNA-seq) (1000 cells per sample) was submitted for single cell barcoding and cDNA synthesis on a 10x Genomics Chromium Controller (10x Genomics), with recovered cell number and sequencing depth for scRNA-seq described in Figure S3. cDNA was sequenced using the NovaSEQ 6000 instrument with an S4 Flow Cell (Illumina). Sequence data was analyzed as follows: Cellranger (2.0.2) count module was used for alignment, filtering, barcode counting and UMI counting of the single cell FASTQs. tSNE coordinates were determined by CellRanger.

#### PrimeFlow RNA Assay

3T12 fibroblasts were infected at an MOI=1.0 with a small-volume inoculation. Inoculum was removed and rinsed-off with PBS after one hour and growth medium was replaced. Cells were harvested 16 hpi and processed for flow cytometry using the PrimeFlow RNA Assay (Thermo Fisher) as described previously (Oko et al., 2019). Briefly, cells were stained with a rabbit antibody against the MHV68 ORF4 protein (vRCA), labeled with a Zenon R-phycoerythrin rabbit IgG label reagent (Life Technologies) per manufacturer’s protocol (Oko et al., 2019). Following cell fixation, cells were then subjected to fluorescent barcoding, labeling distinct experimental conditions with either Ghost Dye Violet 450 (Tonbo Biosciences), Ghost Dye Violet 510 (Tonbo Biosciences) or leaving the cells unlabeled. Labelled samples were then combined into a single tube and subjected to probe hybridization for viral and host RNAs or proteins following the manufacturer’s protocol. Flow cytometric analysis was done on an LSR II instrument (BD Biosciences). Compensation values were initially based on antibody-stained beads (BD Biosciences) and cross-validated using cell samples stained with individual antibody conjugates, with compensation modified as needed post-collection using FlowJo. PrimeFlow probes and antibodies included: i) a type 6 (AlexaFluor750)-mouse actb probe, ii) a type 4 (AF488)-MHV68 ORF18 probe, iii) PE-labelled anti-vRCA antibody, and iv) AF647-conjugated anti-mouse phospho-histone H2AX antibody (clone JBW301, that recognizes phosphorylated serine 139 of H2AX).

#### CyTOF Sample Preparation and Data Acquisition

CyTOF antibodies were obtained from Fluidigm (previously DVS Sciences), provided by the University of Colorado CyTOF Antibody Bank, supported by the Flow Cytometry Shared Resource and the Human Immune Monitoring Shared Resource of the University of Colorado Cancer Center. Samples were FACS purified based on LANA::β-lactamase dependent CCF2-AM cleavage, into either LANAβlac+ or LANAβlac-fractions, transferred to 5mL screw cap tubes, spun down, and stained for viability using 5 μM Cell-ID Cisplatin (Fluidigm) diluted in 1X PBS (Rockland Immunochemicals) for 5 minutes at RT. Samples were then washed, fixed, and each sample labelled with a series of unique palladium isotopic barcodes (Cell-ID 20-Plex Palladium (Pd) Barcoding Kit, Fluidigm). After samples were washed and pooled, cells were pre-treated with 2.4G2 Fc receptor blockade (Tonbo) using a 1:100 dilution of antibody. Cells were incubated with a panel of antibodies, diluted in Maxpar Cell Staining Buffer (Fluidigm), to detect cell surface proteins, washed, then incubated with secondary detection antibodies (anti-FITC, anti-PE, anti-APC, anti-Biotin, and goat anti-rabbit isotopically labelled antibodies) (Kimball et al., 2019). Samples were washed, fixed, and permeabilized (eBioscience Foxp3 / Transcription Factor Staining Buffer Set, Thermo Fisher) before incubation with antibodies to intracellular proteins. The sample was washed, fixed with freshly made 1.6% paraformaldehyde diluted in Maxpar PBS (16% paraformaldehyde, Alfa Aesar), incubated in 0.125nM Iridium intercalator diluted in Maxpar PBS (Fluidigm), and stored at 4°C until acquisition. Samples were washed with 3ml of Maxpar Cell Staining Buffer, washed two times with deionized water (Milli-Q®), resuspended in deionized water (Milli-Q) containing 1:9 EQ Four Element Calibration Beads (Fluidigm), filtered through CellTrics 30um disposable filters (Sysmex-Partec), and counted with a BIORAD TC20 automated cell counter. Data was acquired on the Helios mass cytometer (Fluidigm). To ensure an appropriate flow rate though the Helios, the sample volume was adjusted with the EQ Four Element Calibration Beads (Fluidigm)/deionized water (Milli-Q) to dilute the sample to 1 million cells/mL at an acquired event rate of 400,000-500,000 events/hr.

#### CyTOF sample normalization/debarcoding

Data were normalized with bead-based normalization using NormalizerR2013b_MacOSX and then debarcoded using SingleCellDebarcoderR2013b_ MacOSX, using software downloaded from the Nolan laboratory GitHub page (https://github.com/nolanlab). Normalized and debarcoded data were subjected to traditional Boolean gating in FlowJo, identifying DNA+ events (^191^Iridium (Ir)+ ^193^ Ir+) that were viable (^195^ Platinum(Pt)−), exported for downstream analysis in PhenoGraph.

#### CyTOF Data Analysis

Normalized and debarcoded live DNA+ events were imported into cytofkit, and subjected to the PhenoGraph algorithm. Clustering was done on either 9 cell surface proteins (Figures 1-2) or all 23 proteins (Figure S1). PhenoGraph was run with the following settings: 1) Merge method: “min” (Figures 1, S1) or “ceiling: 10,000 events” (Fig. 2), 2) Transformation method: “cytofAsinh”, 3) Clustering method: “Rphenograph”, and 4) Visualization method: “tSNE”. All other settings were selected by default. Median marker intensity for mock, LANA-, and LANA+ samples was calculated in FlowJo. Statistical significance was tested in GraphPad Prism using an unpaired t test, with statistical significance as identified.

#### PhenoGraph visualization and analysis

PhenoGraph tSNE plots were visualized and customized in the R package “Shiny” (Kimball et al., 2018). In the “Shiny” application cluster color was customized, tSNE plots were colored according to the expression of various cellular markers, and dendrograms were generated. The “cluster cell percentage” .csv file was used for comparing the distribution of events between conditions and determining statistical significance.

#### RT-qPCR

RNA was isolated using the RNeasy Micro Kit (Qiagen) following manufacturer’s specifications. DNA contamination was removed using Turbo-DNAse (ThermoFisher) according to the manufacturer’s specifications. RT-qPCR for Ifnb1 was performed using a primer/probe set (ThermoFisher) and the LightCycler 480 Probes Master (Roche) in 384-well plates on the QuantStudio 7 Flex Instrument (Applied Biosystems, Life Technologies). RNA was reverse transcribed with SuperScript II (Invitrogen) and qPCR was performed using iQ SYBR Green SuperMix (Bio-Rad) in 384-well plates on the QuantStudio 7 Flex instrument (Applied Biosystems, Life Technologies). qRT-PCR results were analyzed using the Pfaffl method, as described by (Pfaffl, 2001), with standardization relative to 18S.

#### Quantification and Statistical Analysis

Data from the 10X Genomics Chromium Controller (10x Genomics) were analyzed with CellRanger, including calculation of cell coordinates for 3D tSNE plots, visualized in Plotly. Unbiased K means clustering was carried out using the DoKmeans function in Seurat. SeqGeq was used for biaxial plotting of scRNAseq data. All flow cytometry data analysis was performed using FlowJo V.10.0.8r1 with flow cytometry data shown as dot plots or averaged for bar graphs. Statistical significance was tested using unpaired student’s t test or one-way ANOVA analysis with Tukey’s multiple comparisons test. Statistical analysis and graphing were performed with GraphPad Prism 7.0d or higher. Gene expression correlation was determined with Spearman’s correlation analysis in GraphPad Prism (8.4.2). CyTOF analysis was done using PhenoGraph as described above or in (Kimball et al., 2018). p values less than 0.05 were considered significant.

### Data and Software Availability

CyTOF data have been deposited to FlowRepository.org and will be made publicly available upon manuscript acceptance. RNA-Seq data have been deposited to NCBI GEO and will be made publicly available upon manuscript acceptance.

### Key Resources Table

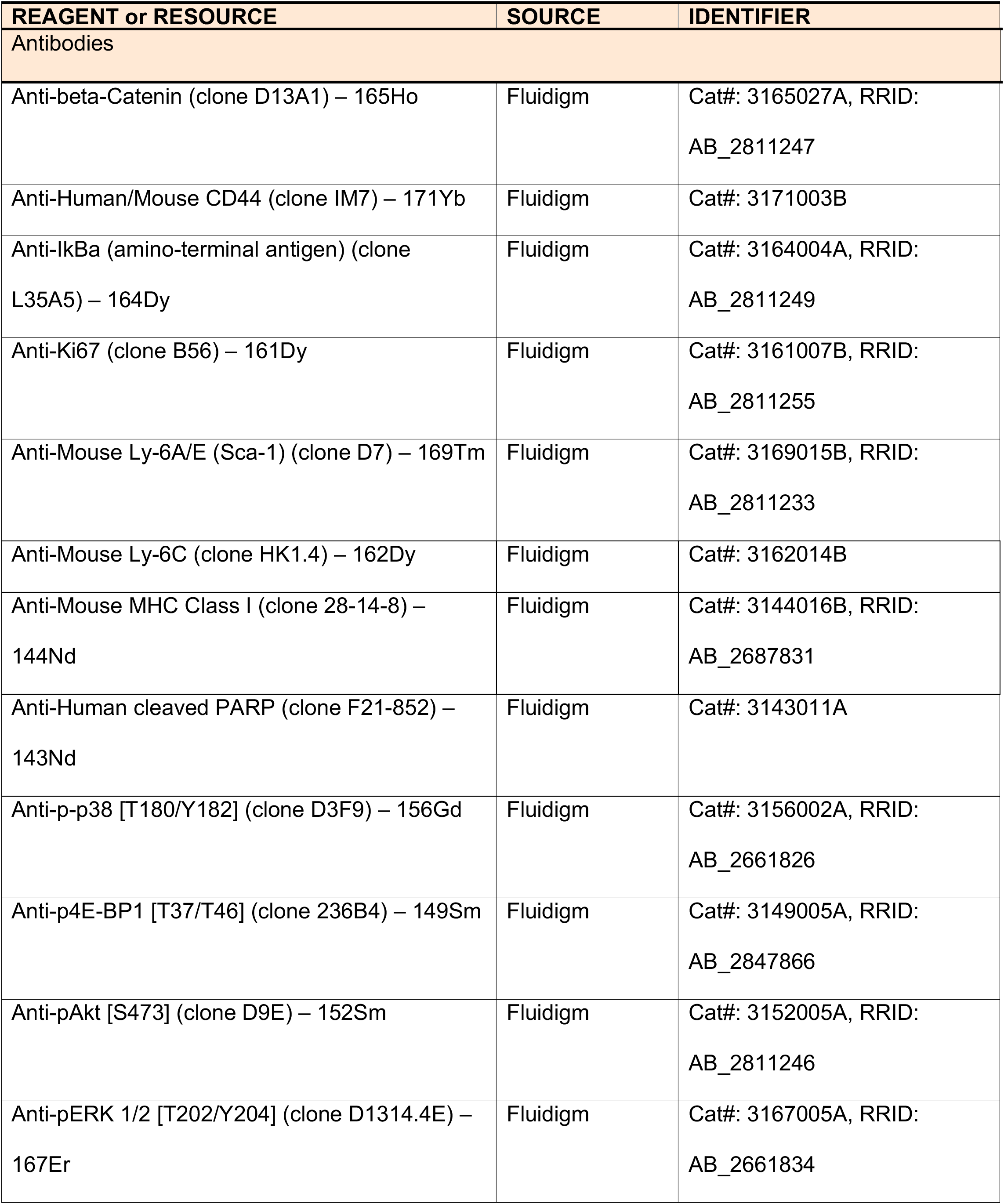
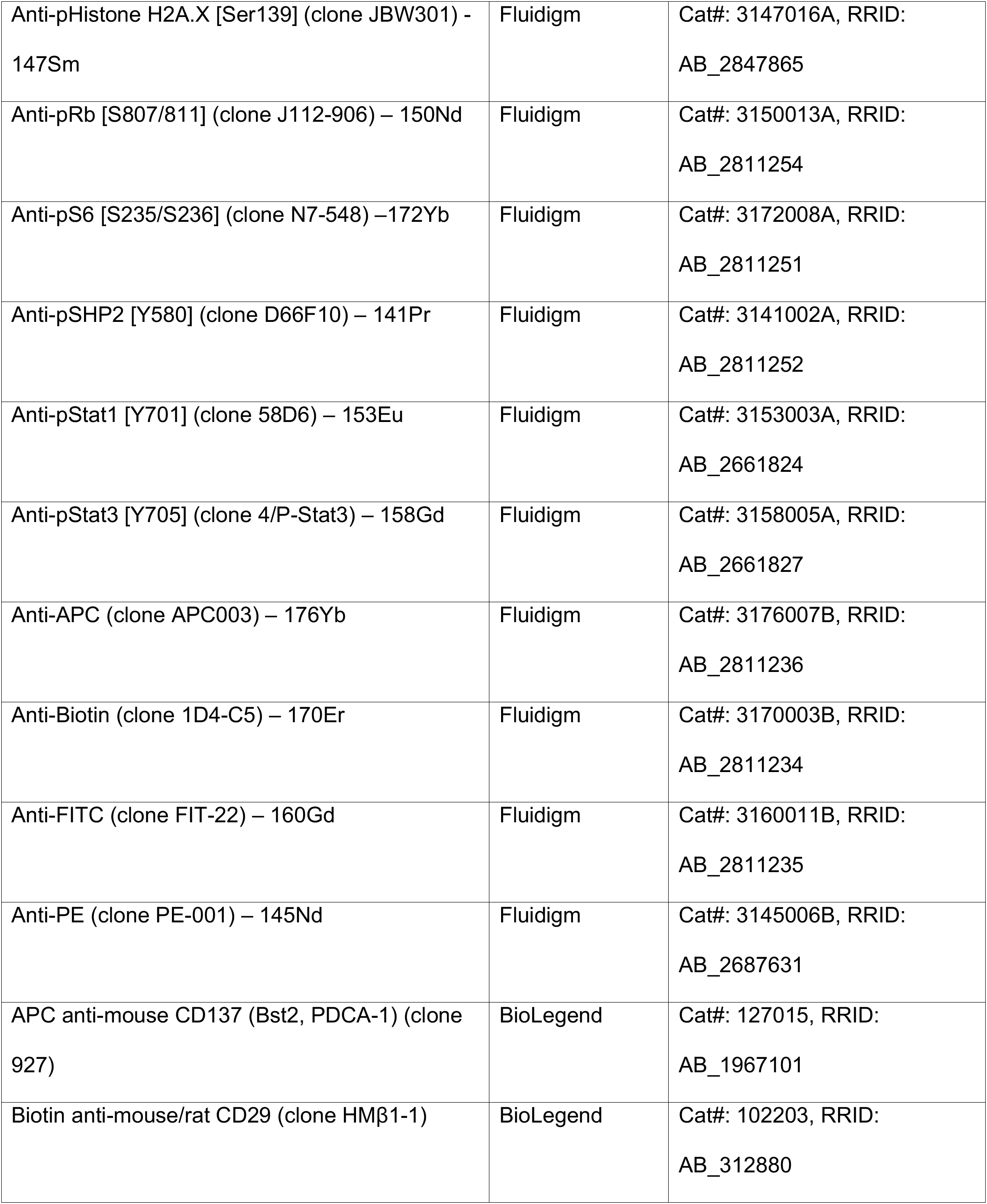
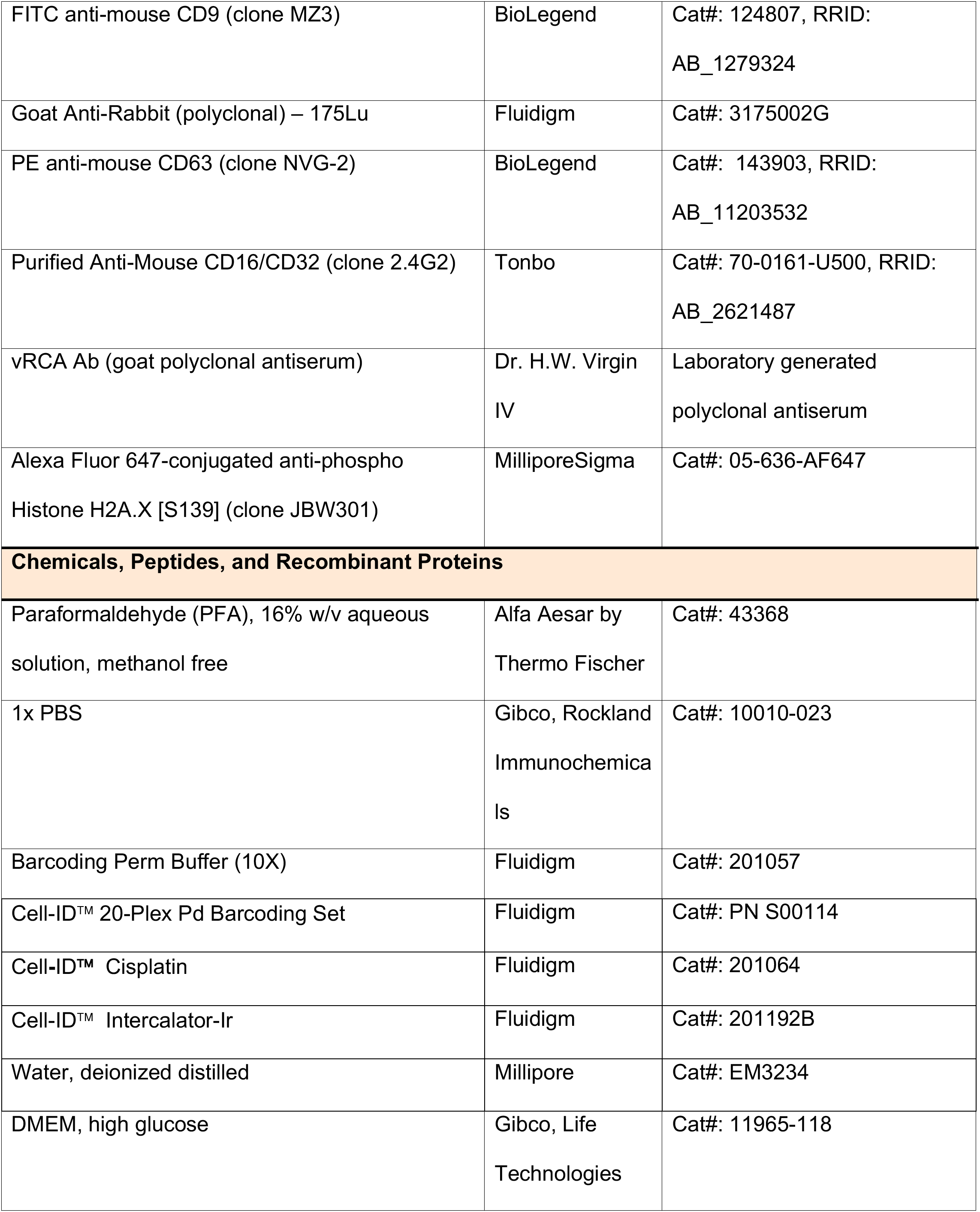
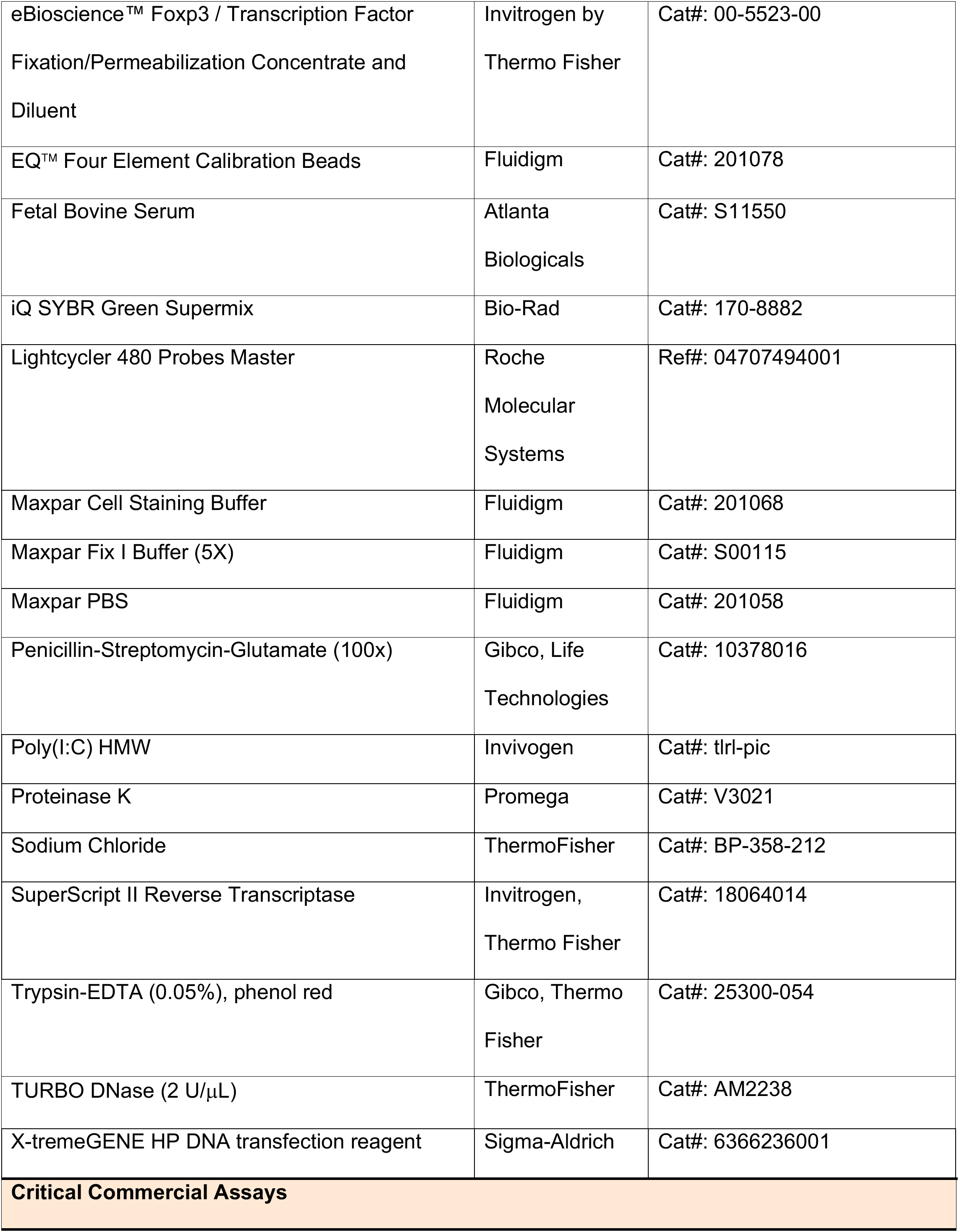
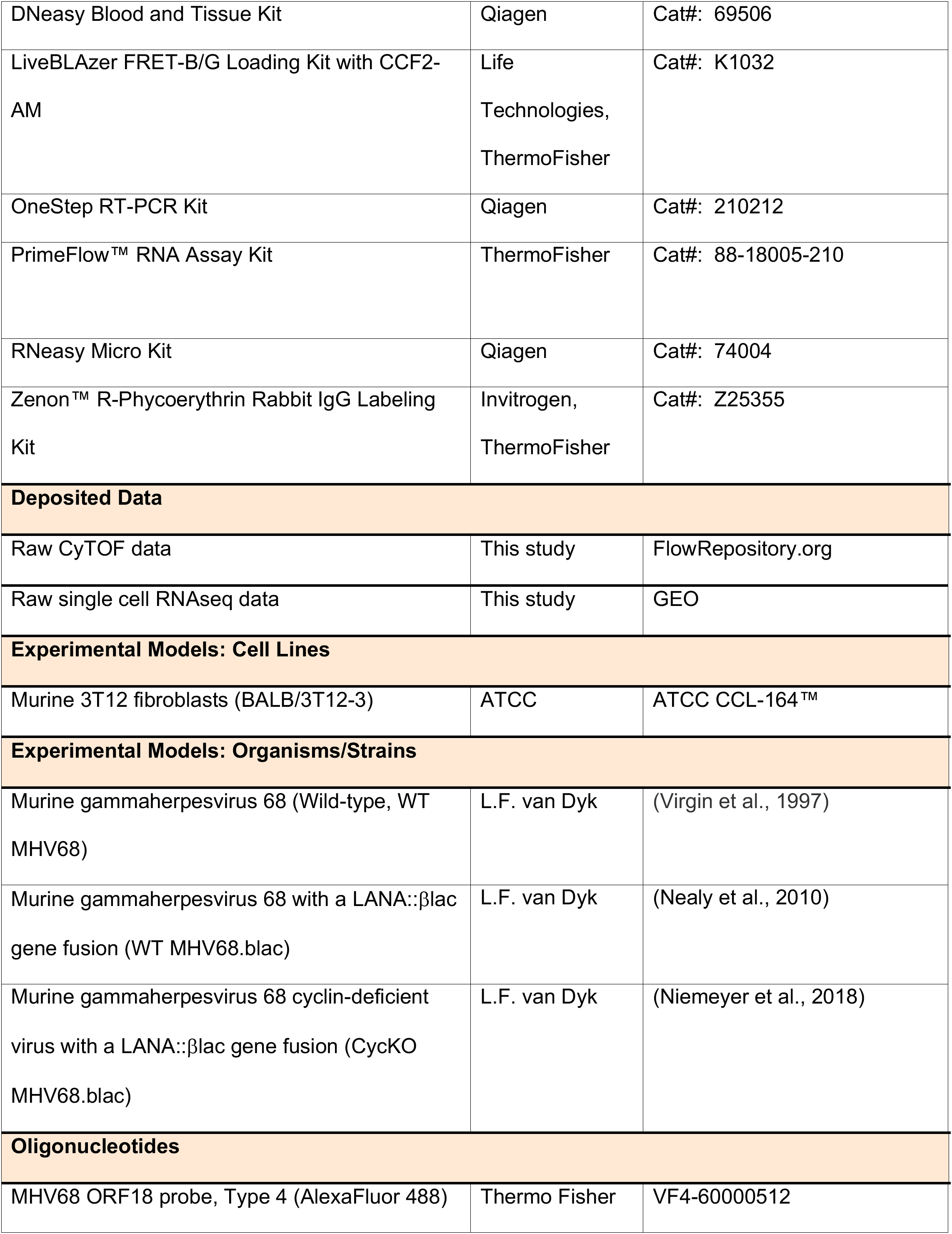
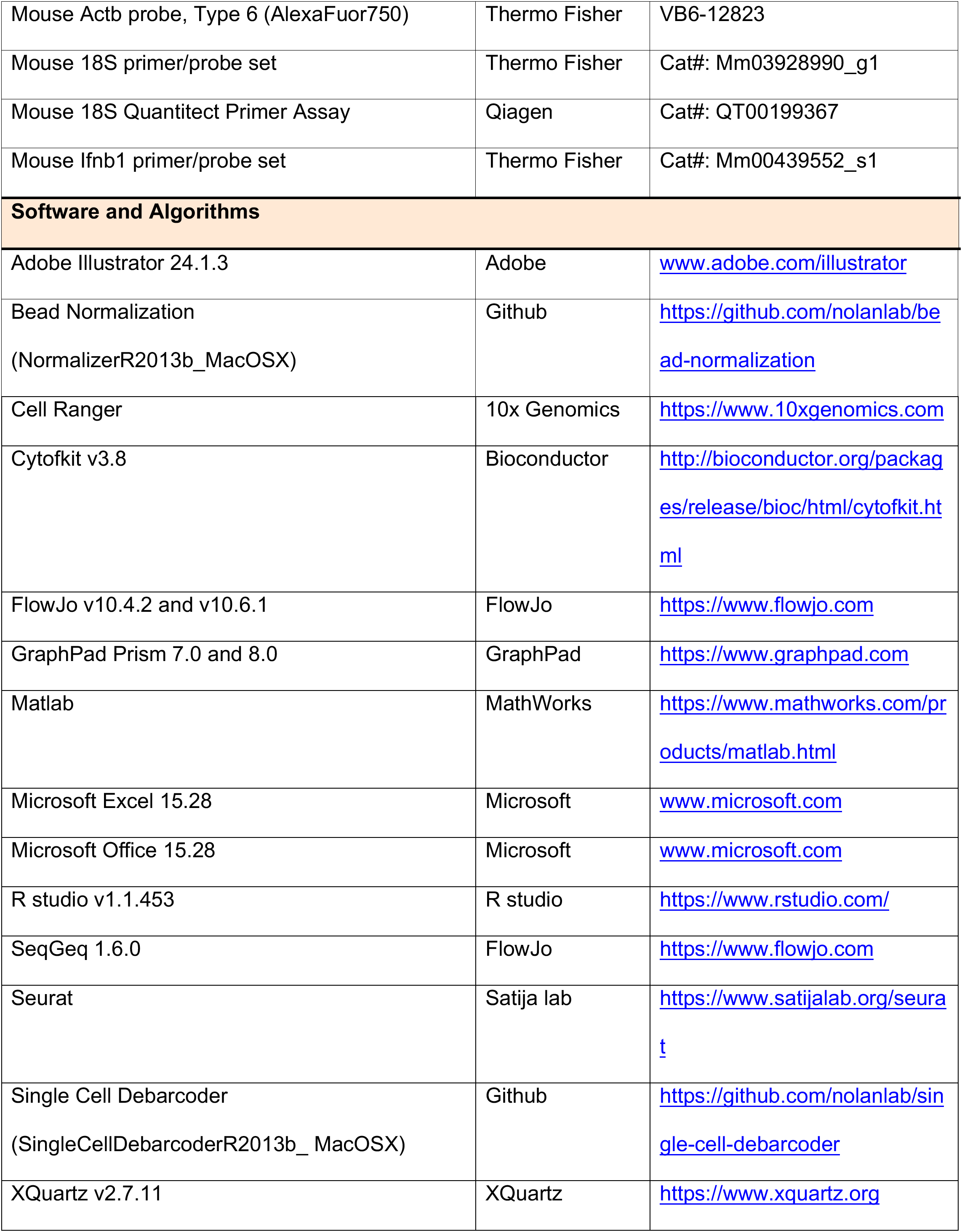

## REFERENCES

Adang, L.A., Parsons, C.H., and Kedes, D.H. (2006). Asynchronous progression through the lytic cascade and variations in intracellular viral loads revealed by high-throughput single-cell analysis of Kaposi’s sarcoma-associated herpesvirus infection. J Virol. 80(20), 10073–10082. Published online 2006/09/29 DOI: 10.1128/JVI.01156-06.

Adler, H., Messerle, M., Wagner, M., and Koszinowski, U.H. (2000). Cloning and mutagenesis of the murine gammaherpesvirus 68 genome as an infectious bacterial artificial chromosome. J Virol. 74(15), 6964–6974. Published online 2000/07/11.

Ahn, J.W., Powell, K.L., Kellam, P., and Alber, D.G. (2002). Gammaherpesvirus lytic gene expression as characterized by DNA array. J Virol. 76(12), 6244–6256. Published online 2002/05/22 DOI: 10.1128/jvi.76.12.6244-6256.2002.

Avraham, R., Haseley, N., Brown, D., Penaranda, C., Jijon, H.B., Trombetta, J.J., Satija, R., Shalek, A.K., Xavier, R.J., Regev, A., et al. (2015). Pathogen Cell-to-Cell Variability Drives Heterogeneity in Host Immune Responses. Cell. 162(6), 1309–1321. Published online 2015/09/08 DOI: 10.1016/j.cell.2015.08.027.

Barton, E., Mandal, P., and Speck, S.H. (2011). Pathogenesis and host control of gammaherpesviruses: lessons from the mouse. Annu Rev Immunol. 29, 351–397. Published online 2011/01/12 DOI: 10.1146/annurev-immunol-072710-081639.

Bhatt, A.P., Wong, J.P., Weinberg, M.S., Host, K.M., Giffin, L.C., Buijnink, J., van Dijk, E., Izumiya, Y., Kung, H.J., Temple, B.R., et al. (2016). A viral kinase mimics S6 kinase to enhance cell proliferation. Proc Natl Acad Sci U S A. 113(28), 7876–7881. Published online 2016/06/28 DOI: 10.1073/pnas.1600587113.

Cesarman, E. (2014). Gammaherpesviruses and lymphoproliferative disorders. Annu Rev Pathol. 9, 349–372. Published online 2013/10/12 DOI: 10.1146/annurev-pathol-012513-104656.

Chang, P.C., Campbell, M., and Robertson, E.S. (2016). Human Oncogenic Herpesvirus and Post-translational Modifications - Phosphorylation and SUMOylation. Front Microbiol. 7, 962. Published online 2016/07/06 DOI: 10.3389/fmicb.2016.00962.

Chaturvedi, S., Klein, J., Vardi, N., Bolovan-Fritts, C., Wolf, M., Du, K., Mlera, L., Calvert, M., Moorman, N.J., Goodrum, F., et al. (2020). A Molecular Mechanism for Probabilistic Bet-hedging and its Role in Viral Latency. bioRxiv. DOI: https://doi.org/10.1101/2020.05.14.096560.

Cheng, B.Y., Zhi, J., Santana, A., Khan, S., Salinas, E., Forrest, J.C., Zheng, Y., Jaggi, S., Leatherwood, J., and Krug, L.T. (2012). Tiled microarray identification of novel viral transcript structures and distinct transcriptional profiles during two modes of productive murine gammaherpesvirus 68 infection. J Virol. 86(8), 4340–4357. Published online 2012/02/10 DOI: 10.1128/JVI.05892-11.

Coleman, H.M., Efstathiou, S., and Stevenson, P.G. (2005). Transcription of the murine gammaherpesvirus 68 ORF73 from promoters in the viral terminal repeats. J Gen Virol. 86(Pt 3), 561–574. Published online 2005/02/22 DOI: 10.1099/vir.0.80565-0.

Covarrubias, S., Gaglia, M.M., Kumar, G.R., Wong, W., Jackson, A.O., and Glaunsinger, B.A. (2011). Coordinated destruction of cellular messages in translation complexes by the gammaherpesvirus host shutoff factor and the mammalian exonuclease Xrn1. PLoS Pathog. 7(10), e1002339. Published online 2011/11/03 DOI: 10.1371/journal.ppat.1002339.

Covarrubias, S., Richner, J.M., Clyde, K., Lee, Y.J., and Glaunsinger, B.A. (2009). Host shutoff is a conserved phenotype of gammaherpesvirus infection and is orchestrated exclusively from the cytoplasm. J Virol. 83(18), 9554–9566. Published online 2009/07/10 DOI: 10.1128/JVI.01051-09.

Damania, B., Jeong, J.H., Bowser, B.S., DeWire, S.M., Staudt, M.R., and Dittmer, D.P. (2004). Comparison of the Rta/Orf50 transactivator proteins of gamma-2-herpesviruses. J Virol. 78(10), 5491–5499. Published online 2004/04/29 DOI: 10.1128/jvi.78.10.5491-5499.2004.

Diebel, K.W., Oko, L.M., Medina, E.M., Niemeyer, B.F., Warren, C.J., Claypool, D.J., Tibbetts, S.A., Cool, C.D., Clambey, E.T., and van Dyk, L.F. (2015). Gammaherpesvirus small noncoding RNAs are bifunctional elements that regulate infection and contribute to virulence in vivo. mBio. 6(1), e01670–01614. Published online 2015/02/19 DOI: 10.1128/mBio.01670-14.

Dong, S., Forrest, J.C., and Liang, X. (2017). Murine Gammaherpesvirus 68: A Small Animal Model for Gammaherpesvirus-Associated Diseases. Adv Exp Med Biol. 1018, 225–236. Published online 2017/10/21 DOI: 10.1007/978-981-10-5765-6_14.

Drayman, N., Patel, P., Vistain, L., and Tay, S. (2019). HSV-1 single-cell analysis reveals the activation of anti-viral and developmental programs in distinct sub-populations. Elife. 8. Published online 2019/05/16 DOI: 10.7554/eLife.46339.

Ebrahimi, B., Dutia, B.M., Roberts, K.L., Garcia-Ramirez, J.J., Dickinson, P., Stewart, J.P., Ghazal, P., Roy, D.J., and Nash, A.A. (2003). Transcriptome profile of murine gammaherpesvirus-68 lytic infection. J Gen Virol. 84(Pt 1), 99–109. Published online 2003/01/21 DOI: 10.1099/vir.0.18639-0.

Forrest, J.C., and Speck, S.H. (2008). Establishment of B-cell lines latently infected with reactivation-competent murine gammaherpesvirus 68 provides evidence for viral alteration of a DNA damage-signaling cascade. J Virol. 82(15), 7688–7699. Published online 2008/05/23 DOI: 10.1128/JVI.02689-07.

Glaunsinger, B.A. (2015). Modulation of the Translational Landscape During Herpesvirus Infection. Annu Rev Virol. 2(1), 311–333. Published online 2016/03/10 DOI: 10.1146/annurev-virology-100114-054839.

Gong, D., Dai, X., Xiao, Y., Du, Y., Chapa, T.J., Johnson, J.R., Li, X., Krogan, N.J., Deng, H., Wu, T.T., et al. (2017). Virus-Like Vesicles of Kaposi’s Sarcoma-Associated Herpesvirus Activate Lytic Replication by Triggering Differentiation Signaling. J Virol. 91(15). Published online 2017/05/19 DOI: 10.1128/JVI.00362-17.

Guenet, J.L. (2005). The mouse genome. Genome Res. 15(12), 1729–1740. Published online 2005/12/13 DOI: 10.1101/gr.3728305.

Hair, J.R., Lyons, P.A., Smith, K.G., and Efstathiou, S. (2007). Control of Rta expression critically determines transcription of viral and cellular genes following gammaherpesvirus infection. J Gen Virol. 88(Pt 6), 1689–1697. Published online 2007/05/09 DOI: 10.1099/vir.0.82548-0.

Hansen, M.M.K., Desai, R.V., Simpson, M.L., and Weinberger, L.S. (2018). Cytoplasmic Amplification of Transcriptional Noise Generates Substantial Cell-to-Cell Variability. Cell Syst. 7(4), 384–397 e386. Published online 2018/09/24 DOI: 10.1016/j.cels.2018.08.002.

Hartenian, E., Gilbertson, S., Federspiel, J.D., Cristea, I.M., and Glaunsinger, B.A. (2020). RNA decay during gammaherpesvirus infection reduces RNA polymerase II occupancy of host promoters but spares viral promoters. PLoS Pathog. 16(2), e1008269. Published online 2020/02/08 DOI: 10.1371/journal.ppat.1008269.

Hartenian, E., and Glaunsinger, B.A. (2019). Feedback to the central dogma: cytoplasmic mRNA decay and transcription are interdependent processes. Crit Rev Biochem Mol Biol. 54(4), 385–398. Published online 2019/10/28 DOI: 10.1080/10409238.2019.1679083.

Hill, E.R., Koganti, S., Zhi, J., Megyola, C., Freeman, A.F., Palendira, U., Tangye, S.G., Farrell, P.J., and Bhaduri-McIntosh, S. (2013). Signal transducer and activator of transcription 3 limits Epstein-Barr virus lytic activation in B lymphocytes. J Virol. 87(21), 11438–11446. Published online 2013/08/24 DOI: 10.1128/JVI.01762-13.

Jang, G.H., Lee, J., Kim, N.Y., Kim, J.H., Yeh, J.Y., Han, M., Ahn, S.K., Kang, H., and Lee, M. (2016). Suppression of lytic replication of Kaposi’s sarcoma-associated herpesvirus by autophagy during initial infection in NIH 3T3 fibroblasts. Arch Virol. 161(3), 595–604. Published online 2015/12/02 DOI: 10.1007/s00705-015-2698-2.

Johnson, L.S., Willert, E.K., and Virgin, H.W. (2010). Redefining the genetics of murine gammaherpesvirus 68 via transcriptome-based annotation. Cell Host Microbe. 7(6), 516–526. Published online 2010/06/15 DOI: 10.1016/j.chom.2010.05.005.

Kapadia, S.B., Molina, H., van Berkel, V., Speck, S.H., and Virgin, H.W.t. (1999). Murine gammaherpesvirus 68 encodes a functional regulator of complement activation. J Virol. 73(9), 7658–7670. Published online 1999/08/10.

Kimball, A.K., Oko, L.M., Bullock, B.L., Nemenoff, R.A., van Dyk, L.F., and Clambey, E.T. (2018). A Beginner’s Guide to Analyzing and Visualizing Mass Cytometry Data. J Immunol. 200(1), 3–22. Published online 2017/12/20 DOI: 10.4049/jimmunol.1701494.

Kimball, A.K., Oko, L.M., Kaspar, R.E., van Dyk, L.F., and Clambey, E.T. (2019). High-Dimensional Characterization of IL-10 Production and IL-10-Dependent Regulation during Primary Gammaherpesvirus Infection. Immunohorizons. 3(3), 94–109. Published online 2019/07/30 DOI: 10.4049/immunohorizons.1800088.

Klasse, P.J. (2015). Molecular determinants of the ratio of inert to infectious virus particles. Prog Mol Biol Transl Sci. 129, 285–326. Published online 2015/01/18 DOI: 10.1016/bs.pmbts.2014.10.012.

Krishnan, H.H., Naranatt, P.P., Smith, M.S., Zeng, L., Bloomer, C., and Chandran, B. (2004). Concurrent expression of latent and a limited number of lytic genes with immune modulation and antiapoptotic function by Kaposi’s sarcoma-associated herpesvirus early during infection of primary endothelial and fibroblast cells and subsequent decline of lytic gene expression. J Virol. 78(7), 3601–3620. Published online 2004/03/16 DOI: 10.1128/jvi.78.7.3601-3620.2004.

Lee, H.R., Amatya, R., and Jung, J.U. (2015). Multi-step regulation of innate immune signaling by Kaposi’s sarcoma-associated herpesvirus. Virus Res. 209, 39–44. Published online 2015/03/23 DOI: 10.1016/j.virusres.2015.03.004.

Levine, J.H., Simonds, E.F., Bendall, S.C., Davis, K.L., Amir el, A.D., Tadmor, M.D., Litvin, O., Fienberg, H.G., Jager, A., Zunder, E.R., et al. (2015). Data-Driven Phenotypic Dissection of AML Reveals Progenitor-like Cells that Correlate with Prognosis. Cell. 162(1), 184–197. Published online 2015/06/23 DOI: 10.1016/j.cell.2015.05.047.

Ma, S.D., Hegde, S., Young, K.H., Sullivan, R., Rajesh, D., Zhou, Y., Jankowska-Gan, E., Burlingham, W.J., Sun, X., Gulley, M.L., et al. (2011). A new model of Epstein-Barr virus infection reveals an important role for early lytic viral protein expression in the development of lymphomas. J Virol. 85(1), 165–177. Published online 2010/10/29 DOI: 10.1128/JVI.01512-10.

Martinez-Guzman, D., Rickabaugh, T., Wu, T.T., Brown, H., Cole, S., Song, M.J., Tong, L., and Sun, R. (2003). Transcription program of murine gammaherpesvirus 68. J Virol. 77(19), 10488–10503. Published online 2003/09/13 DOI: 10.1128/jvi.77.19.10488-10503.2003.

Matthews, N.C., Goodier, M.R., Robey, R.C., Bower, M., and Gotch, F.M. (2011). Killing of Kaposi’s sarcoma-associated herpesvirus-infected fibroblasts during latent infection by activated natural killer cells. Eur J Immunol. 41(7), 1958–1968. Published online 2011/04/22 DOI: 10.1002/eji.201040661.

Messinger, J.E., Dai, J., Stanland, L.J., Price, A.M., and Luftig, M.A. (2019). Identification of Host Biomarkers of Epstein-Barr Virus Latency IIb and Latency III. mBio. 10(4). Published online 2019/07/04 DOI: 10.1128/mBio.01006-19.

Nawandar, D.M., Wang, A., Makielski, K., Lee, D., Ma, S., Barlow, E., Reusch, J., Jiang, R., Wille, C.K., Greenspan, D., et al. (2015). Differentiation-Dependent KLF4 Expression Promotes Lytic Epstein-Barr Virus Infection in Epithelial Cells. PLoS Pathog. 11(10), e1005195. Published online 2015/10/03 DOI: 10.1371/journal.ppat.1005195.

Nealy, M.S., Coleman, C.B., Li, H., and Tibbetts, S.A. (2010). Use of a virus-encoded enzymatic marker reveals that a stable fraction of memory B cells expresses latency-associated nuclear antigen throughout chronic gammaherpesvirus infection. J Virol. 84(15), 7523–7534. Published online 2010/05/21 DOI: 10.1128/JVI.02572-09.

Niemeyer, B.F., Oko, L.M., Medina, E.M., Oldenburg, D.G., White, D.W., Cool, C.D., Clambey, E.T., and van Dyk, L.F. (2018). Host Tumor Suppressor p18(INK4c) Functions as a Potent Cell-Intrinsic Inhibitor of Murine Gammaherpesvirus 68 Reactivation and Pathogenesis. J Virol. 92(6). Published online 2018/01/05 DOI: 10.1128/JVI.01604-17.

O’Grady, T., Feswick, A., Hoffman, B.A., Wang, Y., Medina, E.M., Kara, M., van Dyk, L.F., Flemington, E.K., and Tibbetts, S.A. (2019). Genome-wide Transcript Structure Resolution Reveals Abundant Alternate Isoform Usage from Murine Gammaherpesvirus 68. Cell Rep. 27(13), 3988–4002 e3985. Published online 2019/06/27 DOI: 10.1016/j.celrep.2019.05.086.

Oko, L.M., Kimball, A.K., Kaspar, R.E., Knox, A.N., Coleman, C.B., Rochford, R., Chang, T., Alderete, B., van Dyk, L.F., and Clambey, E.T. (2019). Multidimensional analysis of Gammaherpesvirus RNA expression reveals unexpected heterogeneity of gene expression. PLoS Pathog. 15(6), e1007849. Published online 2019/06/06 DOI: 10.1371/journal.ppat.1007849.

Pavlova, I., Lin, C.Y., and Speck, S.H. (2005). Murine gammaherpesvirus 68 Rta-dependent activation of the gene 57 promoter. Virology. 333(1), 169–179. Published online 2005/02/15 DOI: 10.1016/j.virol.2004.12.021.

Pellett, P.J., and Roizman, B. (2013). Herpesviridae. In: Fields Virology.

Pfaffl, M.W. (2001). A new mathematical model for relative quantification in real-time RT-PCR. Nucleic Acids Res. 29(9), e45. Published online 2001/05/09 DOI: 10.1093/nar/29.9.e45.

Purushothaman, P., Thakker, S., and Verma, S.C. (2015). Transcriptome analysis of Kaposi’s sarcoma-associated herpesvirus during de novo primary infection of human B and endothelial cells. J Virol. 89(6), 3093–3111. Published online 2015/01/02 DOI: 10.1128/JVI.02507-14.

Rochford, R., Lutzke, M.L., Alfinito, R.S., Clavo, A., and Cardin, R.D. (2001). Kinetics of murine gammaherpesvirus 68 gene expression following infection of murine cells in culture and in mice. J Virol. 75(11), 4955–4963. Published online 2001/05/03 DOI: 10.1128/JVI.75.11.4955-4963.2001.

Ruiz, J.C., Hunter, O.V., and Conrad, N.K. (2019). Kaposi’s sarcoma-associated herpesvirus ORF57 protein protects viral transcripts from specific nuclear RNA decay pathways by preventing hMTR4 recruitment. PLoS Pathog. 15(2), e1007596. Published online 2019/02/21 DOI: 10.1371/journal.ppat.1007596.

Sen, N., Mukherjee, G., Sen, A., Bendall, S.C., Sung, P., Nolan, G.P., and Arvin, A.M. (2014). Single-cell mass cytometry analysis of human tonsil T cell remodeling by varicella zoster virus. Cell Rep. 8(2), 633–645. Published online 2014/07/22 DOI: 10.1016/j.celrep.2014.06.024.

Shaffer, S.M., Dunagin, M.C., Torborg, S.R., Torre, E.A., Emert, B., Krepler, C., Beqiri, M., Sproesser, K., Brafford, P.A., Xiao, M., et al. (2017). Rare cell variability and drug-induced reprogramming as a mode of cancer drug resistance. Nature. 546(7658), 431–435. Published online 2017/06/14 DOI: 10.1038/nature22794.

Shnayder, M., Nachshon, A., Krishna, B., Poole, E., Boshkov, A., Binyamin, A., Maza, I., Sinclair, J., Schwartz, M., and Stern-Ginossar, N. (2018). Defining the Transcriptional Landscape during Cytomegalovirus Latency with Single-Cell RNA Sequencing. mBio. 9(2). Published online 2018/03/15 DOI: 10.1128/mBio.00013-18.

Smiley, J.R. (2004). Herpes simplex virus virion host shutoff protein: immune evasion mediated by a viral RNase? J Virol. 78(3), 1063–1068. Published online 2004/01/15 DOI: 10.1128/jvi.78.3.1063-1068.2004.

Suarez, A.L., and van Dyk, L.F. (2008). Endothelial cells support persistent gammaherpesvirus 68 infection. PLoS Pathog. 4(9), e1000152. Published online 2008/09/13 DOI: 10.1371/journal.ppat.1000152.

Tarakanova, V.L., Leung-Pineda, V., Hwang, S., Yang, C.W., Matatall, K., Basson, M., Sun, R., Piwnica-Worms, H., Sleckman, B.P., and Virgin, H.W.t. (2007). Gamma-herpesvirus kinase actively initiates a DNA damage response by inducing phosphorylation of H2AX to foster viral replication. Cell Host Microbe. 1(4), 275–286. Published online 2007/11/17 DOI: 10.1016/j.chom.2007.05.008.

van Dyk, L.F., Virgin, H.W.t., and Speck, S.H. (2000). The murine gammaherpesvirus 68 v-cyclin is a critical regulator of reactivation from latency. J Virol. 74(16), 7451–7461. Published online 2000/07/25 DOI: 10.1128/jvi.74.16.7451-7461.2000.

Van Skike, N.D., Minkah, N.K., Hogan, C.H., Wu, G., Benziger, P.T., Oldenburg, D.G., Kara, M., Kim-Holzapfel, D.M., White, D.W., Tibbetts, S.A., et al. (2018). Viral FGARAT ORF75A promotes early events in lytic infection and gammaherpesvirus pathogenesis in mice. PLoS Pathog. 14(2), e1006843. Published online 2018/02/02 DOI: 10.1371/journal.ppat.1006843.

Virgin, H.W.t., Latreille, P., Wamsley, P., Hallsworth, K., Weck, K.E., Dal Canto, A.J., and Speck, S.H. (1997). Complete sequence and genomic analysis of murine gammaherpesvirus 68. J Virol. 71(8), 5894–5904. Published online 1997/08/01.

Virgin, H.W.t., Presti, R.M., Li, X.Y., Liu, C., and Speck, S.H. (1999). Three distinct regions of the murine gammaherpesvirus 68 genome are transcriptionally active in latently infected mice. J Virol. 73(3), 2321–2332. Published online 1999/02/11.

Waterboer, T., Rahaus, M., and Wolff, M.H. (2002). Varicella-zoster virus (VZV) mediates a delayed host shutoff independent of open reading frame (ORF) 17 expression. Virus Genes. 24(1), 49–56. Published online 2002/04/04 DOI: 10.1023/a:1014086004141.

Weitzman, M.D., and Fradet-Turcotte, A. (2018). Virus DNA Replication and the Host DNA Damage Response. Annu Rev Virol. 5(1), 141–164. Published online 2018/07/12 DOI: 10.1146/annurev-virology-092917-043534.

Wu, T.T., Usherwood, E.J., Stewart, J.P., Nash, A.A., and Sun, R. (2000). Rta of murine gammaherpesvirus 68 reactivates the complete lytic cycle from latency. J Virol. 74(8), 3659–3667. Published online 2000/03/23 DOI: 10.1128/jvi.74.8.3659-3667.2000.

Wyler, E., Franke, V., Menegatti, J., Kocks, C., Boltengagen, A., Praktiknjo, S., Walch-Ruckheim, B., Bosse, J., Rajewsky, N., Grasser, F., et al. (2019). Single-cell RNA-sequencing of herpes simplex virus 1-infected cells connects NRF2 activation to an antiviral program. Nat Commun. 10(1), 4878. Published online 2019/10/28 DOI: 10.1038/s41467-019-12894-z.

Zamora, M.R. (2011). DNA viruses (CMV, EBV, and the herpesviruses). Semin Respir Crit Care Med. 32(4), 454–470. Published online 2011/08/23 DOI: 10.1055/s-0031-1283285.

Zhang, K., Lv, D.W., and Li, R. (2019). Conserved Herpesvirus Protein Kinases Target SAMHD1 to Facilitate Virus Replication. Cell Rep. 28(2), 449–459 e445. Published online 2019/07/11 DOI: 10.1016/j.celrep.2019.04.020.

