## Supplemental Figures S1-7, Supplemental Tables S1-3 for "Redefining De Novo Gammaherpesvirus Infection Through High-Dimensional, Single-Cell Analysis of Virus and Host"

### Supplemental Figures and Tables for "Redefining De Novo Gammaherpesvirus Infection Through High-Dimensional, Single-Cell Analysis of Virus and Host"

Jennifer N. Berger[1], Bridget Sanford[2], Abigail K. Kimball[3], Lauren M. Oko[1], Rachael E. Kaspar[3], Brian F. Niemeyer[1], Kenneth L. Jones[4], Eric T. Clambey[3]*, Linda F. van Dyk[1,5]*

**Affiliations**
[1] Department of Immunology and Microbiology, University of Colorado Anschutz Medical Campus | Aurora, CO, 80045, USA
[2] Department of Pediatric Oncology, University of Colorado Anschutz Medical Campus | Aurora, CO, 80045, USA
[3] Department of Anesthesiology, University of Colorado Anschutz Medical Campus | Aurora, CO, 80045, USA
[4] Department of Cell Biology, University of Oklahoma Health Sciences Center | Oklahoma City, OK, 73104, USA
[5] Lead Contact

**Supplemental Figures, Figure Legends and Tables:**
- Supplemental Figures and Figure legends S1-7
- Supplemental Tables S1-3

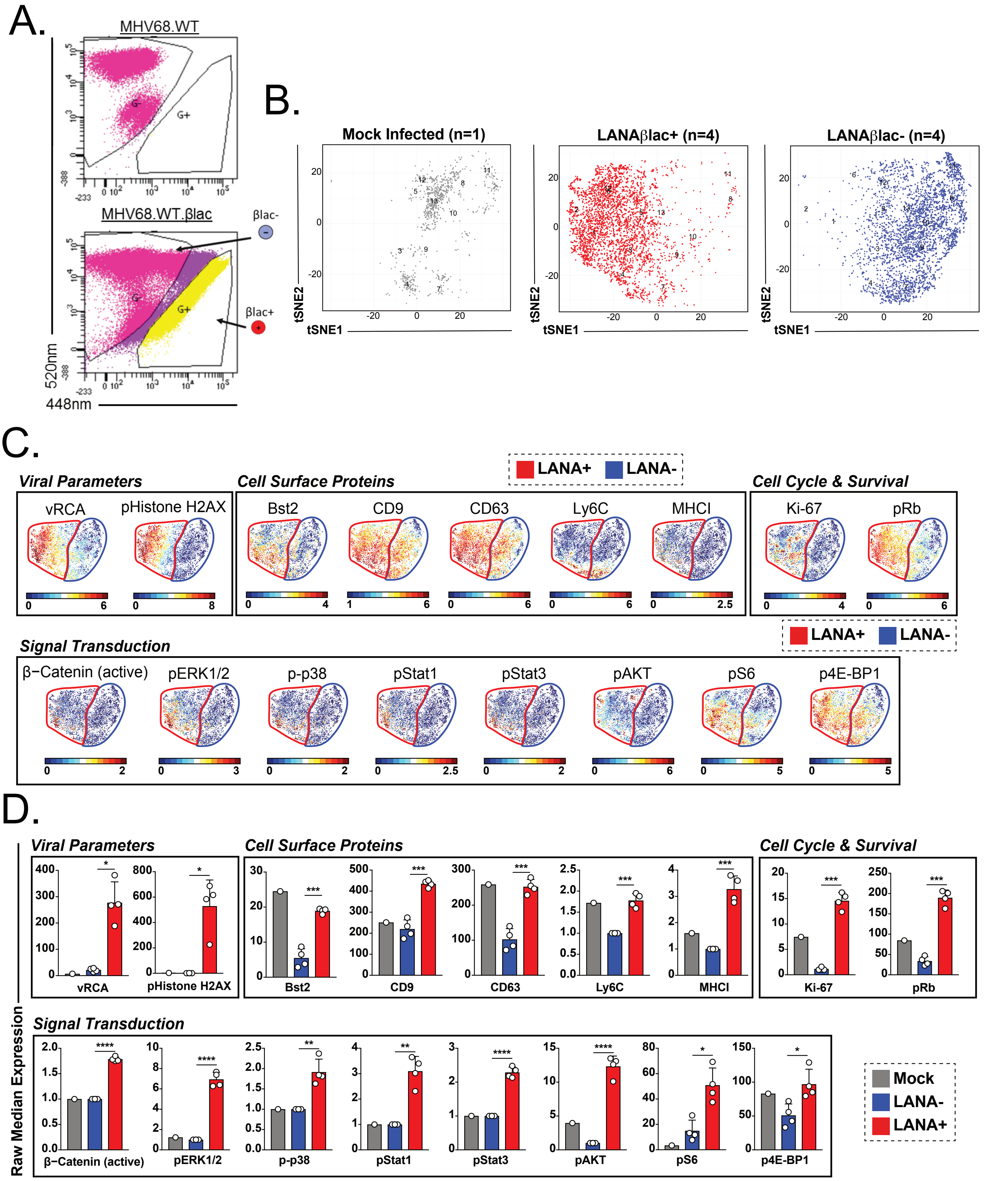


Figure S1
Berger et al

**Figure S1. CyTOF analysis identifies distinct protein expression profiles for LANA+ virally-infected cells.** High-dimensional single-cell protein analysis of MHV68 infection by CyTOF as depicted in Figure 1. (A) Flow cytometry gates for sorting LANAβlac- and LANAβlac+ cells. Infection with WT MHV68 not containing the β-lactamase marker (MHV68.WT, top) serves as a critical control for the definition of LANA+ cells after MHV68 infection containing the β-lactamase marker (MHV68.WTblac, bottom). (B) Cellular distribution of mock, LANA- and LANA+ cells subjected to CyTOF analysis and visualized by tSNE-based dimensionality reduction identified 14 cell clusters. (C) tSNE-based visualization of protein expression, comparing LANA+ (red boundary) and LANA- (blue boundary) for the indicated protein targets. Ruler indicates range of expression for each protein target, with values representing arcsinh (x/5) where x is raw expression value. (D) Protein expression profiles that are statistically significantly different between LANA+ (red) and LANA- (blue) cells. Data depict mean ± SEM of raw median with individual symbols indicating values from independent samples. Proteins are arranged by functional categories. Live, DNA+ cells ($^{191}$Ir+ $^{193}$Ir+ $^{195}$Pt-) were imported into PhenoGraph with 9,162 events total (1,018 events from each sample; n=1, mock, n=4 each for LANA+ and LANA- samples) clustered on 23 protein markers (see STAR methods). All samples were analyzed for statistical significance using unpaired t tests, corrected for multiple comparisons using the Holm-Sidak method, with statistical significance denoted as *p<0.05, ** p<0.01, ***p<0.001.

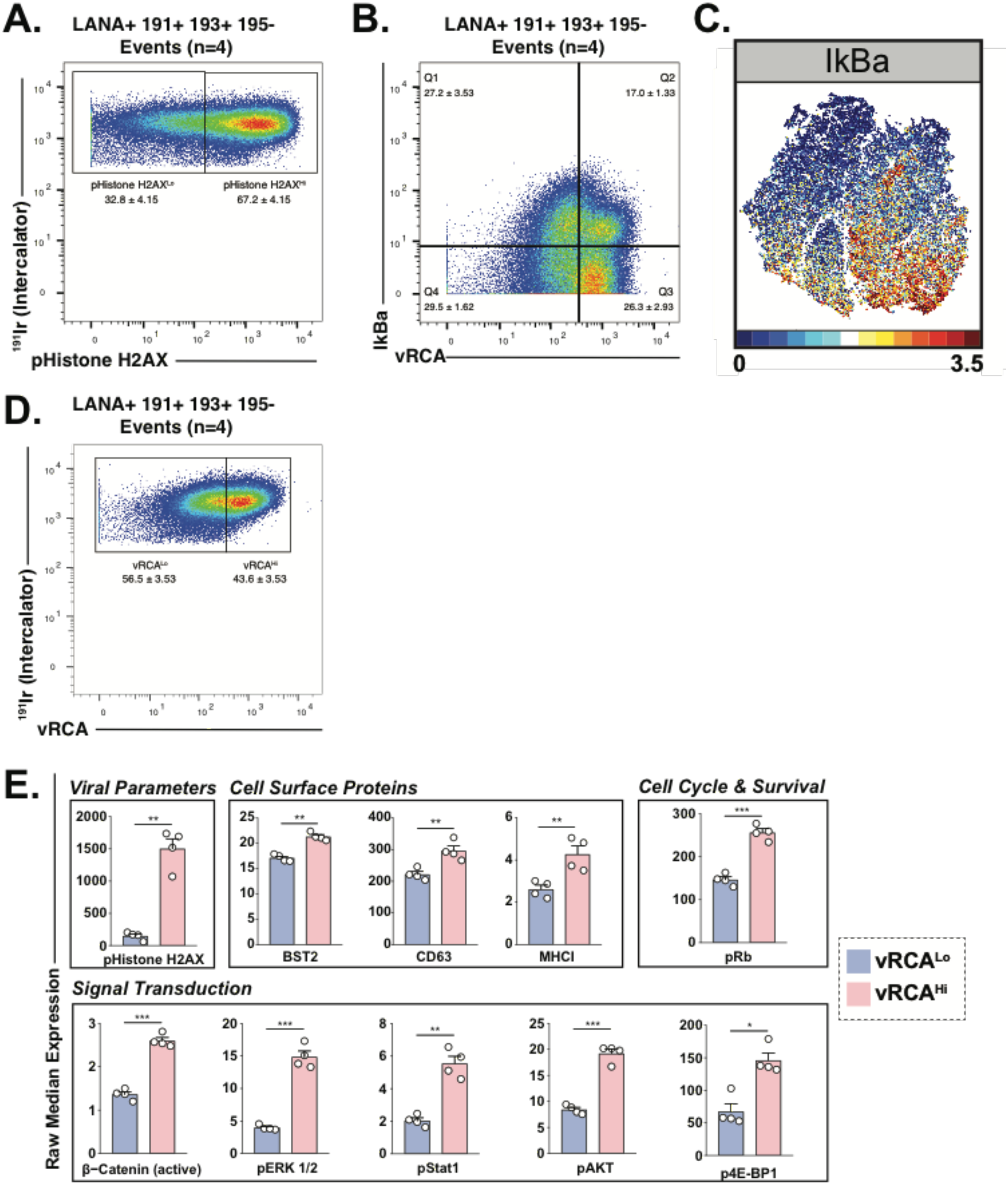


Figure S2
Berger et al

**Figure S2. CyTOF analysis of LANA+ cells and the use of pH2AX and vRCA as indicators of infection.** High-dimensional single-cell protein analysis of MHV68 infection by CyTOF as depicted in Figure 1. (A) LANA+ live, DNA+ events ($^{191}Ir$+ $^{193}Ir$+ $^{195}Pt$-) were stratified into $pH2AX^{hi}$ and $pH2AX^{lo}$ subsets for detailed analysis in Figure 2. (B-C) IkBa expression differs based on vRCA expression, with $vRCA^{hi}$ cells characterized by modestly reduced IkBa levels by (B) biaxial plot or (C) tSNE-based visualization of IkBa (parallel cell layout as in Figure 2C, with ruler indicating expression range and values representing arcsinh (x/5) where x is raw expression value). (D-E) LANA+ live, DNA+ events were stratified based on vRCA expression into $vRCA^{hi}$ and $vRCA^{lo}$ subsets and compared for raw median expression. Data depict protein expression that was statistically significantly different between $vRCA^{hi}$ and $vRCA^{lo}$ subsets. Data depict mean ± SEM with individual symbols indicating values from independent samples (n=4 samples/group). All samples were analyzed for statistical significance using unpaired t tests, corrected for multiple comparisons using the Holm-Sidak method, with statistical significance denoted as *$p<0.05$, **$p<0.01$, ***$p<0.001$. Protein markers quantified by CyTOF that were not differentially expressed are not shown.

A.

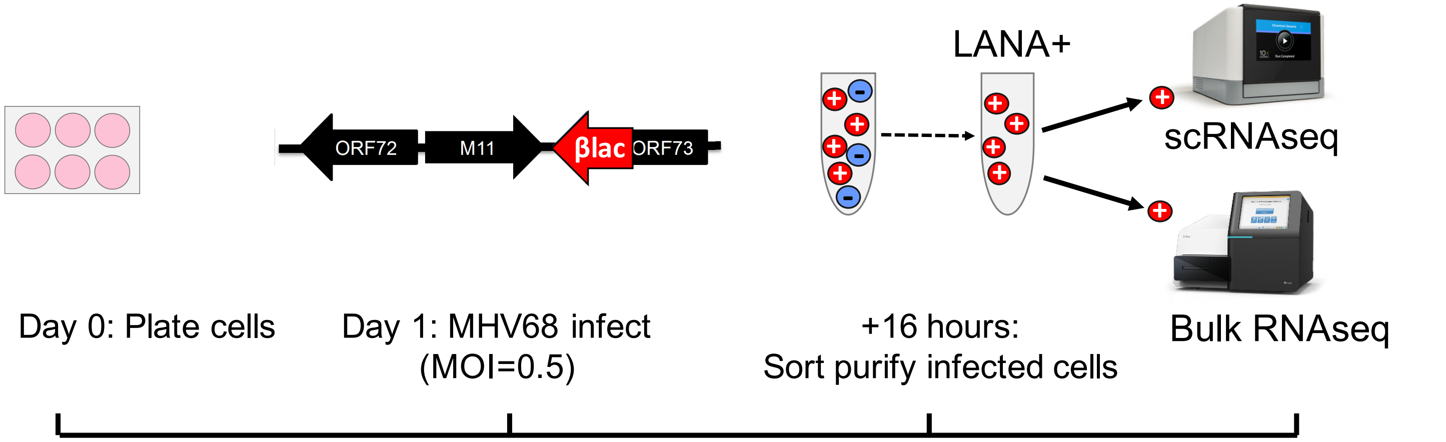


B.

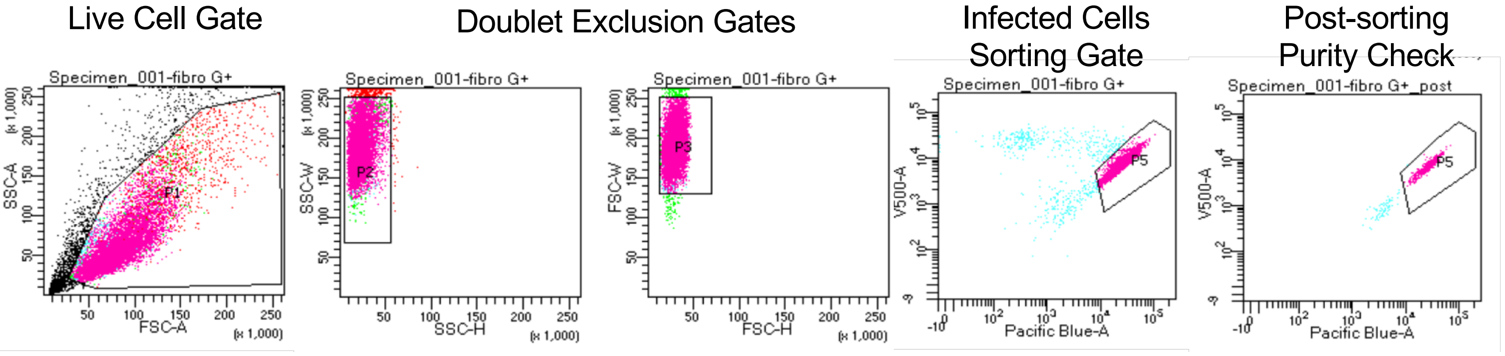


C.

| | |
|---|---|
| **Cell #** | 1,605 |
| **Post-Normalization Mean Reads /Cell** | 94,810 |
| **Median Genes/Cell** | 1,148 |

D.

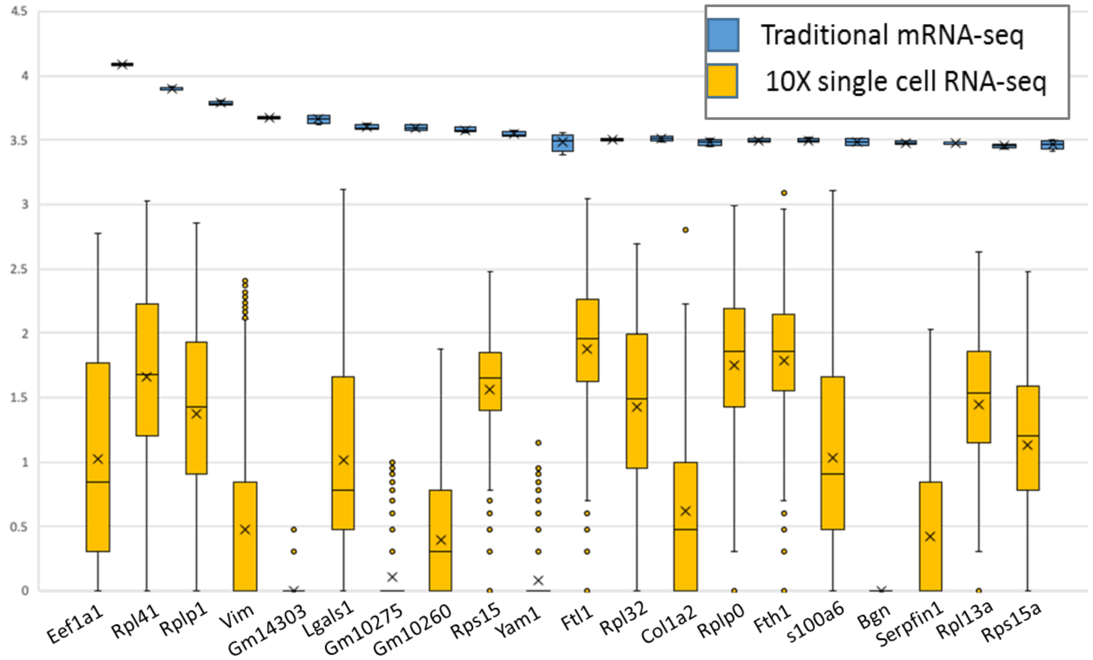


Figure S3
Berger et al

**Figure S3. Comparison of traditional versus single cell RNA-seq analysis of LANA+ MHV68 infected fibroblasts.** (A) Schematic of MHV68 infection for scRNAseq and bulk RNAseq. 3T12 cells were inoculated with MHV68 that contains the LANA::β-lactamase fusion gene, MOI=0.5 for 1 hour, and harvested at 16 hpi cells by staining with β-lactamase substrate. (B) Infected cells were purified by fluorescence activated cell sorting (FACS) for β-lactamase positivity. (C) The number of cells, mean reads per cell, and median genes per cell from scRNA-seq. (D) The top 20 detected genes in traditional bulk RNAseq (blue) and scRNA-seq (gold) were plotted as log2 expression values from original FPKM and UMI data, respectively. Box boundaries represent the first and the third quartiles, x = mean, horizontal bar = median, whiskers = SD. N = 4 samples for bulk RNA-seq; N = 1605 cells for scRNA-seq. Comparison demonstrates notable variation in expression between single cells, even in the most highly expressed genes.

| | WT MHV68 | CycKO MHV68 |
|---|---|---|
| **All cells** | | |
| Cell number | 859 | 746 |
| Sum of virus UMIs | 4,399,078 | 3,721,678 |
| Sum of host UMIs | 9,309,084 | 7,833,273 |
| Ratio of virus:host | 0.473 | 0.475 |
| **Virus-biased cells** | | |
| Cell number | 630 (73.3%) | 546 (73.2%) |
| Sum of virus UMIs | 4,104,588 | 3,509,875 |
| Sum of host UMIs | 2,569,468 | 2,030,618 |
| **Host-biased cells** | | |
| Cell number | 229 (26.7%) | 200 (26.8%) |
| Sum of virus UMIs | 289,972 | 216,321 |
| Sum of host UMIs | 6,737,129 | 5,805,142 |

**Table S1. Cell and gene distribution in scRNA-seq dataset.** Cells infected with WT or CycKO MHV68 have comparable representation among the total cells, virus-biased or host-biased cells, with similar distribution of RNAs mapped to either virus or host genomes. Virus-biased and host-biased cells defined in Figure 6E-F.

| Ensembl ID | Common ID | p-value | WT UMIs | CycKO UMIs | Ratio WT/CycKO | Fold change: WT/CycKO |
|---|---|---|---|---|---|---|
| GAMMAHV.ORF68 | GAMMAHV.ORF68 | 1.87E-61 | 4.759022 | 2.096515 | 2.269968 | 1.182672 |
| GAMMAHV.ORF37 | GAMMAHV.ORF37 | 1.13E-37 | 9.486612 | 4.489276 | 2.113172 | 1.07941 |
| GAMMAHV.ORF75c | GAMMAHV.ORF75c | 2.96E-35 | 4.98603 | 2.715818 | 1.835922 | 0.876505 |
| GAMMAHV.ORF59 | GAMMAHV.ORF59 | 2.73E-24 | 2.087311 | 1.217158 | 1.714905 | 0.778129 |
| GAMMAHV.ORF17 | GAMMAHV.ORF17 | 4.67E-22 | 345.7311 | 307.2346 | 1.1253 | 0.17031 |
| GAMMAHV.ORF55 | GAMMAHV.ORF55 | 1.32E-17 | 2.841676 | 1.72252 | 1.64972 | 0.722221 |
| ENSMUSG00000097971 | Gm26917 | 6.8E-13 | 1.076834 | 0.514745 | 2.091973 | 1.064865 |
| ENSMUSG00000060143 | Gm10076 | 8.42E-13 | 4.317811 | 2.077748 | 2.078121 | 1.05528 |
| GAMMAHV.ORF6 | GAMMAHV.ORF6 | 1.65E-11 | 2.157159 | 1.364611 | 1.580787 | 0.660643 |
| ENSMUSG00000044609 | Gm9294 | 1.72E-10 | 5.159488 | 3.175603 | 1.624727 | 0.700197 |
| GAMMAHV.ORF48 | GAMMAHV.ORF48 | 6.67E-08 | 10.65308 | 5.75067 | 1.852494 | 0.889469 |
| ENSMUSG00000072692 | Rpl37rt | 2.78E-06 | 6.565774 | 8.630027 | 0.760806 | -0.3944 |
| GAMMAHV.ORF66 | GAMMAHV.ORF66 | 2.85E-06 | 49.42026 | 59.05898 | 0.836795 | -0.25705 |
| ENSMUSG00000069117 | Gm10260 | 4.79E-06 | 5.672875 | 3.05496 | 1.856939 | 0.892927 |
| GAMMAHV.M9 | GAMMAHV.M9 | 5.39E-06 | 723.4144 | 587.7909 | 1.230734 | 0.299519 |
| ENSMUSG00000098178 | Gm42418 | 7.31E-06 | 2.745052 | 2.227882 | 1.232135 | 0.301161 |
| GAMMAHV.ORF72 | GAMMAHV.ORF72 | 0.000022 | 6.253783 | 29.75737 | 0.210159 | -2.25045 |

**Table S2. There are few differences between WT and CycKO MHV68 lytic infection as defined by scRNA-seq.** Identification of all genes that have expression differences of statistical significance ($p \leq 0.0001$ between wild-type (WT) and cyclin knockout (CycKO) MHV68 infected cells), and for which either WT or CycKO infected cells had $\geq 1$ UMI. Average UMIs for each gene in cells infected with either WT or CycKO viruses are shown in columns four and five respectively. The UMI ratio of WT/CycKO was calculated for each gene (column six). Fold change of the difference in gene expression between WT and CycKO viruses for each gene by taking the $\log_2$ of the expression ratio (last column).

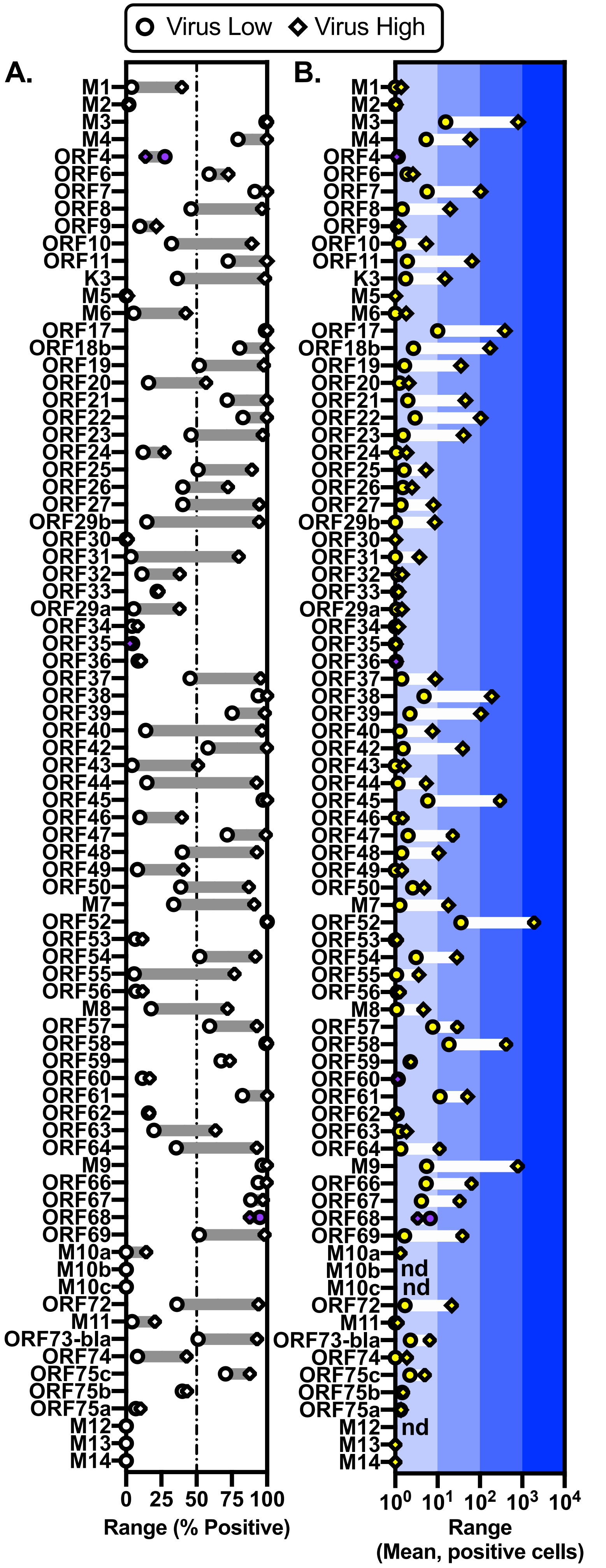


Figure S4
Berger et al

**Figure S4. Analysis of virus gene expression in virus low and virus high cells.** (A) The percent of positive cells for each viral gene (≥1 viral UMI/cell) for virus low (<500 viral UMIs per cell, depicted by circles) and virus high (>500 viral UMIs per cell, depicted by diamonds) groups was calculated. Range between virus low and high groups per gene is indicated with shaded gray bars. Genes with a greater percentage of positive cells in the virus low group are denoted by purple shading of shapes (ORF4, ORF35, ORF68). Dashed line indicates 50% gene expression for reference. (B) Mean UMIs for positive cells (≥1 viral UMI/cell) was calculated for virus low and virus high cells, with mean calculated from only cells that expressed the indicated gene. Range in mean expression depicted by white bars. Background shades of blue indicate 10-fold differences in gene expression. nd, not detected. Virus high (n=1347 cells) and virus low (n=258 cells) groups defined in Figure 4.

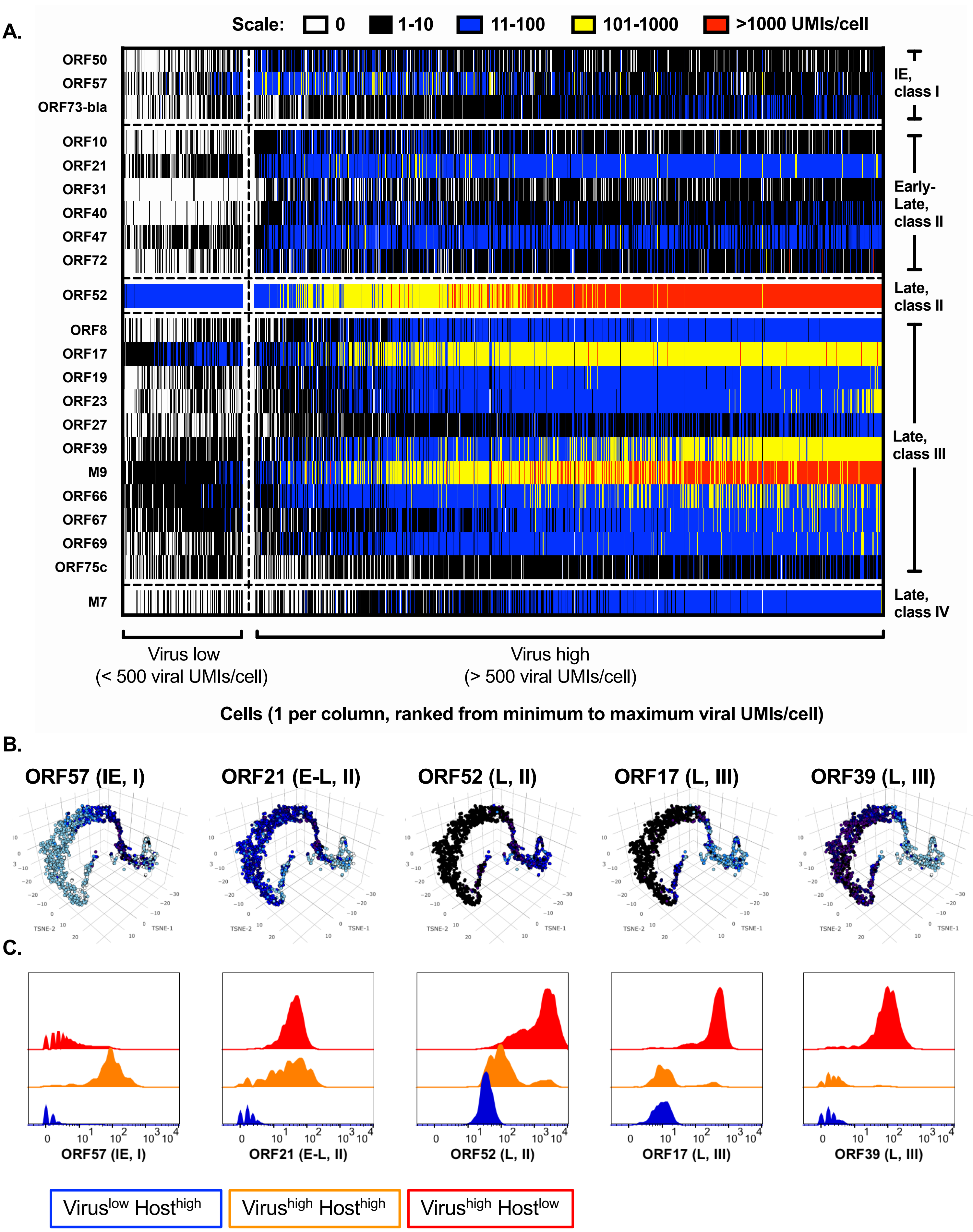


Figure S5
Berger et al

**Figure S5. The frequency and magnitude of viral gene expression arranged according to kinetic class and IE/E/L paradigm.** Immediate early (IE), early (E), early-late (E-L), and late (L) genes were selected based on transcriptional and kinetic class. (A) UMIs were plotted per cell and organized left to right from lowest to highest viral UMIs. Each row depicts expression for a different viral gene. Each column depicts expression within an individual cell. Data depict all LANA+ cells (n=1605 cells). Data correspond to that shown in Figure 5, with viral gene order reorganized based on which category each viral gene falls into. (B) 3D tSNE-based visualization of 5 representative viral genes, comparable to data shown in Figure 6B. Panels A and B depict all LANA+ cells (n=1605 cells). (C) Histogram overlay comparison of viral gene expression as a function of cell subset, with cells stratified into Virus$^{low}$ (blue), Virus$^{high}$ Host$^{high}$ (orange), and Virus$^{high}$ Host$^{low}$ (red) cell subsets as defined in Figure 6K. Kinetic class ("class") for each viral gene is indicated as a roman numeral behind IE, E, E-L or L gene distinction.

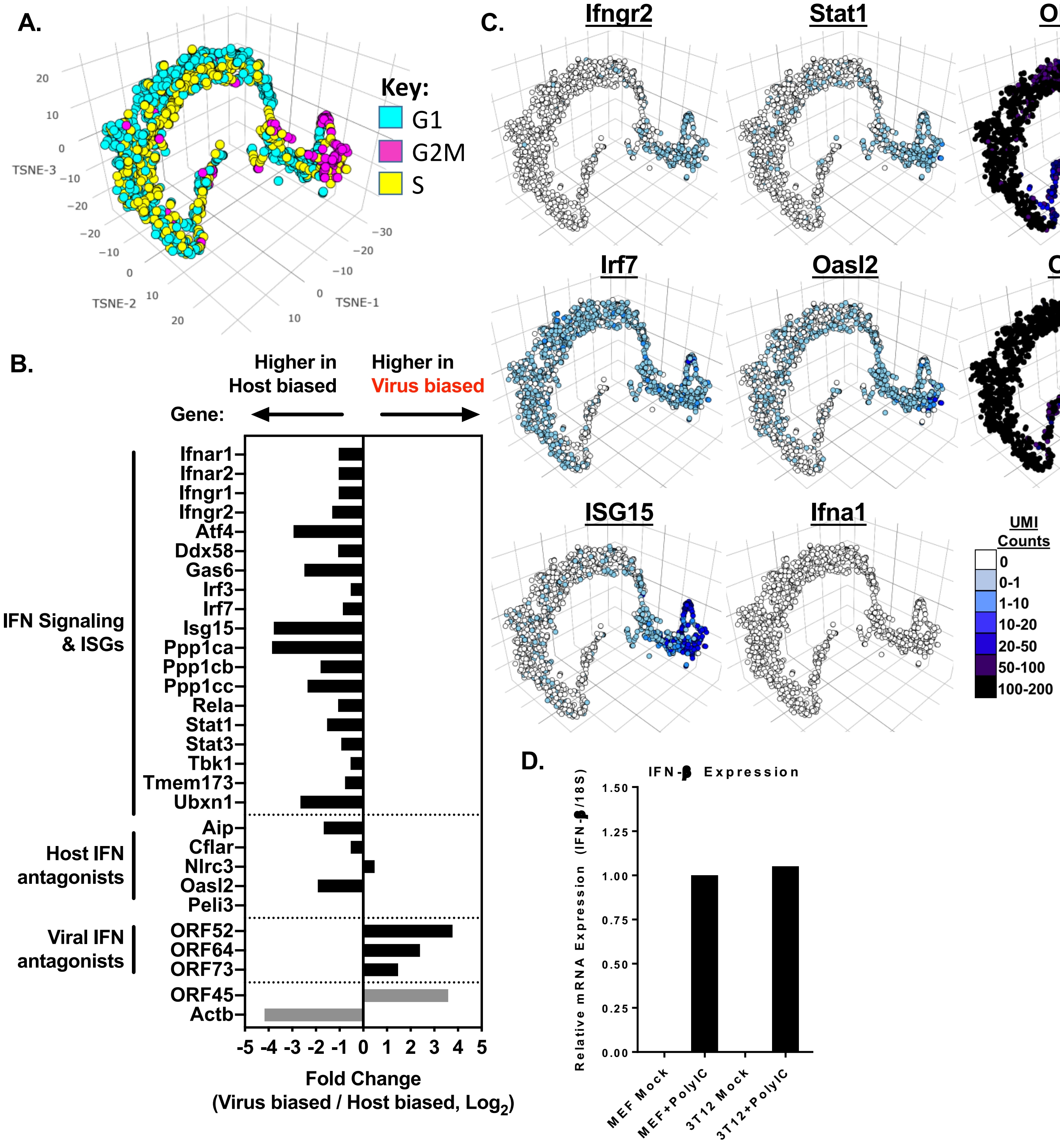


Figure S6
Berger et al

**Figure S6. Potential host cell effects on gene expression during virus infection.** (A) Cell cycle stage was assigned to each cell (Seurat), color coded, and overlaid on 3D tSNE visualization from CellRanger analysis of scRNA-seq data in parallel orientation to that presented in Figure 6. Cell cycle stage is indicated by the color of each cell (legend on right). (B) Interferon association of host-biased and virus-biased genes are graphed based on fold change (calculated from log2 of the ratio of UMIs of virus-biased vs host-biased for the genes indicated) representative of three groups: interferon (IFN) signaling and interferon stimulated genes (ISGs) (top), host IFN antagonists (middle), and viral IFN antagonists (bottom). ORF45 is a viral gene with higher expression in virus-biased cells, in contrast to host Actb, with higher expression in host-biased cells. (C) Representative tSNE plots for genes involved in innate immune signaling, with UMI counts indicated for each cell by color key. (D) Fibroblasts can express IFN-β in response to a synthetic dsRNA ligand. Murine embryonic fibroblasts (MEFs) and 3T12s were transfected with Poly (IC) (+PolyIC) or left untreated (mock). 24 hours after transfection, interferon-β (IFN-β) expression was analyzed with RT-qPCR. Mean expression of six technical replicates is shown, N=1.

| Gene Name | Mean UMIs Host-biased Group | Mean UMIs Virus-biased Group |
|---|---|---|
| Actb | 80.55 | 3.52 |
| ORF45 | 26.75 | 332.01 |
| Isg15 | 16.25 | 0.26 |
| Ubxn1 | 6.80 | 0.24 |
| Irf7 | 5.31 | 2.46 |
| Stat1 | 2.20 | 0.11 |
| Ddx58 | 1.34 | 0.12 |
| Stat3 | 1.12 | 0.12 |
| Tmem173 | 0.75 | 0.02 |
| Tbk1 | 0.69 | 0.17 |
| Irf3 | 0.53 | 0.06 |
| Ifngr2 | 1.56 | 0.03 |
| Ifnar2 | 1.15 | 0.05 |
| Ifngr1 | 1.10 | 0.02 |
| Ifnar1 | 0.48 | 0.12 |
| Ifna1 | 0.00 | 0.00 |
| ORF52 | 155.32 | 2137.36 |
| ORF73 | 1.74 | 6.59 |
| ORF64 | 1.36 | 11.41 |
| Oasl2 | 4.82 | 0.54 |
| Aip | 2.63 | 0.14 |
| Cflar | 0.55 | 0.07 |
| Nlrc3 | 0.06 | 0.47 |
| Peli3 | 0.03 | 0.00 |

**Table S3. IFN associated genes are expressed within the host-biased cell subset.** Mean UMIs for genes involved in interferon signaling in the host-biased group (second column) compared to the virus-biased group (last column). ORF45, ORF52, ORF73 and ORF64 are viral genes, with all other genes encoded by the host. Host-biased and virus-biased populations defined in Figure 6.

Figure S7
Berger et al

| A. All cells (n=1605 cells) | |
|---|---|
| B. Virus$^{Low}$ (n=258) | C. Virus$^{High}$ (n=1347) |
| D. Virus$^{High}$ Host$^{High}$ (n=178) | E. Virus$^{High}$ Host$^{Low}$ (n=1169) |

A.

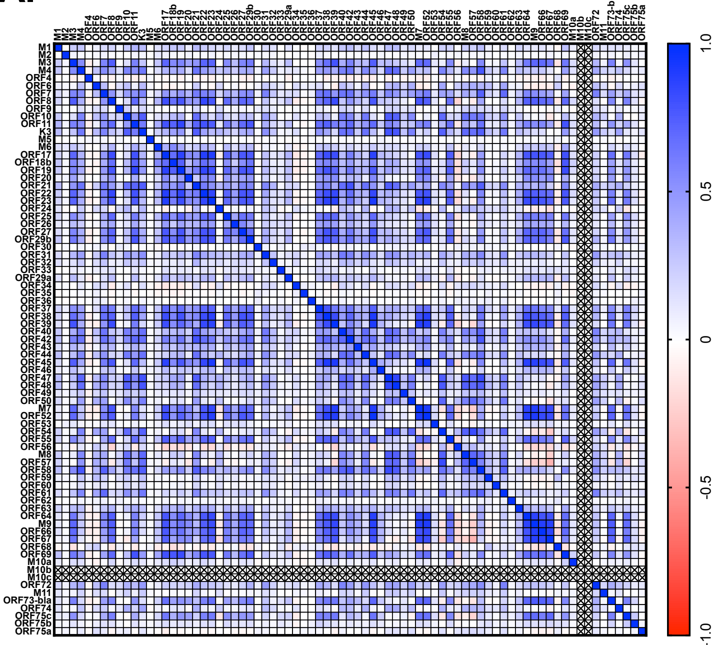


B. C. D. E.

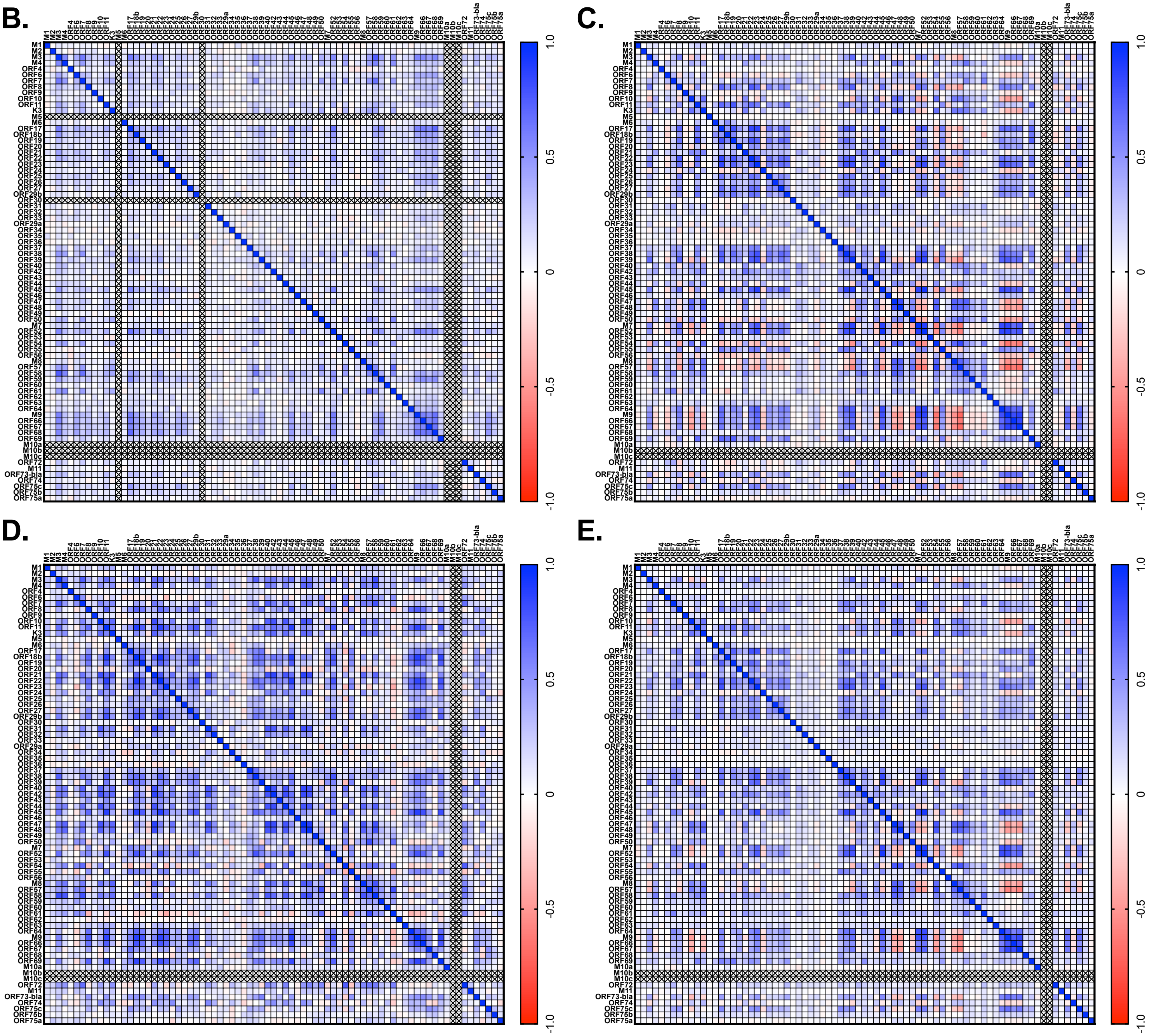

**Figure S7. Gene expression analysis reveals positive and negative correlations in MHV68 transcription at the single-cell level.** The inter-relationship of viral gene expression was interrogated using scRNA-seq data from MHV68-infected LANA+ cells purified as in Figure 1A. Correlation matrix of viral genes among (A) all cells, (B) $Virus^{Low}$ cells (<500 viral UMIs/cell), (C) $Virus^{High}$ cells (>500 viral UMIs/cell), (D) $Virus^{High}$ $Host^{High}$ cells and (E) $Virus^{High}$ $Host^{Low}$ cells (as defined in Figure 6), depicting Spearman correlation coefficient, rs, for each gene pair according to the indicated heatmap. Positive correlation is indicated by blue, negative correlation indicated by red. Correlation matrix shows all viral genes except for M10b and M10c (X indicates gene for which there were no values) and M12, M13 and M14 with extremely low values. These data contain and expand upon data presented in Figure 7.
